# Water Kefir as a Model for Aggregative Development of Microbial Multicellularity within a Multispecies Biofilm

**DOI:** 10.64898/2026.10.09.757837

**Authors:** Pia Bethge, Berenike Schmalfuß, Samuel Ropert, Florian Schattenberg, Gorkhmaz Abbaszade, Susann Müller, Thomas Henle, Yael Politi, Thorsten Mascher

## Abstract

Water kefir is a traditional fermented beverage and a non-dairy alternative to milk kefir. Water kefir grains are cauliflower-like aggregates, ranging in size from 1–20 mm. They are composed of an exopolysaccharide matrix derived from the fermentation substrate sucrose. The microbial community on and within the grains is a complex, highly stable multicellular aggregate typically consisting of 10–15 core species that primarily belong to three major groups: lactic acid bacteria, acetic acid bacteria, and yeasts. Remarkably, the structural organisation, stability, and growth of kefir grains cannot be readily explained by their constituent species alone, and attempts to reconstitute them from the isolated species have so far been unsuccessful. Here, we combined a comprehensive structural investigation of the macro- and micromorphology of water kefir grains with culture-dependent and -independent analyses of its microbial biodiversity. The dynamics and interactions within this complex microbial community were studied by combining flow cytometry, fluorescence-activated cell sorting, and amplicon sequencing of selected subpopulations. Furthermore, the growth of water kefir grains was spatiotemporally resolved and quantitatively characterised, thereby unravelling an emergent multicellular growth function, documented for the first time. Our combined evidence provides an unprecedented understanding of a highly structured and metabolically synergistic multicellular microbial aggregate and lets us propose water kefir as a model system for the aggregative development of microbial multicellularity from multiple species. It should lay the foundation for future mechanistic studies aimed at understanding how multispecies microbial communities self-organise into stable, spatially structured multicellular aggregates.

**IMPORTANCE:** Water kefir is an example of aggregative microbial multicellularity. Its grains consist of a highly resilient symbiotic community of bacteria and yeasts. Although the microbial and chemical compositions of kefir grains are well understood, the macroscopic mechanisms of growth and multiplication, as well as the internal grain architecture, are poorly understood. Here, we demonstrate that kefir grains grow by fragmenting existing grains once they reach a critical, destabilising size. This unique and multicellular mode of division is a function of the kefir grain architecture, which resembles a microbial ‘planet’ with a dense but uninhabited core, with cell division only occurring in the outer shell of the grains. Water kefir grains provide a fascinating microbial model system for studying aggregative multicellularity and the relationships between multicellular structure and function.

## INTRODUCTION

Multicellularity – the organisation of multiple cells into a coordinated, functionally integrated unit – represents one of the most significant transitions in the history of life on Earth [1,2]. It is one of a small number of major evolutionary events in which the unit of selection shifted from individual organisms to collectives [1]. Multicellularity has evolved at least 25 times independently across the tree of life, including in animals, plants, fungi, algae, and bacteria, suggesting that the selective pressures favouring cellular cooperation are both potent and widespread [3,4]. The recurrent emergence of multicellularity points to fundamental benefits such as division of labour, increased size conferring predation resistance, more efficient resource exploitation, and protection from environmental stresses [5,6].

Within the context of multicellularity, a critical conceptual distinction is made between clonal and aggregative origins [7]. In clonal multicellularity, cells remain associated following division, so that the resulting group consists of genetically identical or near-identical individuals. This genetic homogeneity suppresses within-group conflict and facilitates the evolution of altruistic or cooperative traits, including programmed cell death and irreversible differentiation [6,7]. Animal development is the paradigmatic example, but clonal multicellularity is also found in volvocine green algae, providing powerful comparative models for studying the stepwise transition from unicellularity to complex organisation [8]. In aggregative multicellularity, by contrast, free-living cells of potentially diverse genotypes are recruited into a common structure, generating the possibility of cheating and within-group competition [7,9]. The social amoeba *Dictyostelium discoideum* and the fruiting-body-forming myxobacteria exemplify this strategy and have become central study systems for understanding how cooperation is maintained despite relatedness asymmetries [9,10].

Bacteria were long considered predominantly unicellular, yet it is now well established that multicellular organisation is widespread and evolutionarily ancient in the prokaryotic domain [11]. Bacterial multicellularity encompasses a remarkable diversity of forms: the programmed development of spore-forming chains in *Bacillus*, the differentiation of heterocysts in filamentous cyanobacteria, the coordinated predatory swarms and fruiting bodies of myxobacteria, and the structured communities of biofilms [11,12]. These systems exhibit hallmark features of multicellularity, including cell differentiation, intercellular communication, coordinated collective behaviour, and, in some cases, a germ-soma distinction [12,13]. Importantly, bacterial multicellularity is subject to the same evolutionary logic as its eukaryotic counterparts: kin selection and high relatedness within clonal groups promote cooperative investment, while mechanisms that suppress defection maintain group-level adaptations [10,14]. The study of bacterial multicellularity, therefore, not only illuminates the ecology and pathogenesis of medically and environmentally important organisms but also offers tractable experimental models for dissecting the genetic and mechanistic foundations of one of evolution’s most transformative innovations [15].

Aggregative multicellularity, in which free-living cells are recruited into a shared structure without obligate clonal descent, finds some of its most compelling microbial expressions in biofilms [16]. In single-species biofilms, planktonic cells transition to a surface-attached, matrix-enclosed lifestyle through a developmental programme involving initial reversible attachment, irreversible anchoring, microcolony formation, and eventual dispersal [17]. Within these structures, cells are far from homogeneous: steep gradients of oxygen, nutrients, and signalling molecules generate distinct physiological niches that drive phenotypic differentiation, such that cells at the colony periphery, interior, and base adopt markedly different gene expression profiles and functional roles [18]. Properties such as division of labour, intercellular signalling, emergent collective properties such as programmed cell death, and vulnerability to within-group defection establish single-species biofilms as bona fide, albeit rudimentary, instances of aggregative multicellularity [11,12].

The conceptual and biological complexity increases considerably when aggregative multicellularity involves more than one microbial species, a condition that is the rule rather than the exception in natural and host-associated environments [19]. Multi-species microbial consortia exhibit properties that are not merely additive combinations of their constituent members but emergent consequences of inter-species interactions encompassing metabolic cross-feeding, physical co-aggregation, horizontal gene transfer, and interspecific signalling [14,20]. The oral cavity provides the most extensively characterised example of multi-species aggregative multicellularity in the form of dental plaque, a structured biofilm community harbouring several hundred bacterial species whose spatial organisation is non-random and whose assembly follows reproducible successional trajectories [20–22]. Beyond the oral cavity, multi-species aggregative multicellularity is increasingly recognised in other body sites and environmental niches. Chronic wound biofilms typically harbour polymicrobial communities dominated by *Staphylococcus aureus* and *Pseudomonas aeruginosa* [23,24]. In the gut, spatially structured mucosal biofilms composed of dozens of bacterial species mediate host–microbiome interactions critical for immune homeostasis and nutrient absorption [25]. Environmental examples include syntrophic consortia of sulphate-reducing bacteria and methanogenic archaea, which are physically associated in structured aggregates and rely on interspecies transfer of hydrogen or formate to sustain their collective energy metabolism – a relationship so tightly coupled that neither partner can sustain growth in isolation under the prevailing conditions [26]. Taken together, these examples demonstrate that aggregative multicellularity in microbial systems spans a continuum from the relatively simple cooperative dynamics of single-species biofilms to the elaborate multi-species consortia.

Water kefir grains are a remarkable, but so far underappreciated, example of aggregative multicellularity involving multiple microbial species. Water kefir, also referred to as ’sugary kefir’, is a traditional fermented beverage and a non-dairy, vegan alternative to milk kefir [27]. It is typically made by dissolving sucrose in water, adding dried fruit, such as figs, as a nitrogen source, and inoculating the solution with water kefir grains, which contain a variety of microorganisms that drive fermentation [28–31]. The fermentation typically proceeds at room temperature for 1–3 days, depending on the desired product characteristics. The main metabolites produced from the substrate are lactic acid, acetic acid, gluconic acid, ethanol, and carbon dioxide. In addition, other organic acids, complex alcohols, and volatile organic compounds, such as esters, ethers, and fatty acids, are produced and contribute to the beverage’s unique flavour [28,30,32]. After each fermentation, the grains are separated from the fermented liquid by sieving and then rinsed before being reused in the next fermentation [33]. The origin of water kefir grains is not documented, but two sources are most often cited: the pads of Mexican *Opuntia* cacti or the Caucasus [34,35].

Remarkably, the *de novo* synthesis of water kefir grains is currently not possible. Cultures can therefore be propagated only through the natural growth and multiplication of pre-existing grains [35,36]. Due to the lack of defined starter cultures, the inability to synthesise grains, product variations, and limited shelf life, significant industrial production has so far been prevented [35,37]. Therefore, water kefir is primarily produced and distributed by private households.

Water kefir grains consist of an exopolysaccharide (EPS) matrix, primarily made of dextran produced from sucrose by lactic acid bacteria (LAB). This matrix is synthesised via extracellular β-glucancrases that catalyse the transfer of glucose units, forming α-1,6-linked backbones with various branching patterns [38]. EPS production is common among water kefir-associated genera such as *Leuconostoc* and (*Liquori-*) *Lactobacillus* [39–41]. The resulting grain surface appears smooth to irregular and cauliflower-like. Individual grains range in size from 1–20 mm, have a soft, elastic, and gel-like texture, and are typically translucent to milky white [35,42].

The microbial community of the grains includes mainly LAB, acetic acid bacteria (AAB), and yeasts. Across studies, 10–15 core species are typically found, with common bacterial genera including *Liquorilactobacillus*, *Lacticaseibacillus, Lentilactobacillus, Bifidobacterium, Zymomonas, Acetobacter,* and *Gluconobacter*, and yeasts from *Saccharomyces, Zygotorulaspora, Pichia,* and *Brettanomyces/Dekkera* [28,29,32,43–45]. Despite overlaps, each culture tends to harbour a unique but stable species composition.

Previous studies have shown that the grain surface is heterogeneously colonised by bacterial and yeast cells [46–48], and that its interior is sparsely populated and contains substantial amounts of dextran, as well as empty areas, which were classified as gas inclusions [49]. However, a comprehensive understanding of the architecture and biogeography of water kefir grains as a whole is lacking. Likewise, it is not known to what extent the population dynamics are affected by the growth cycle. In fact, even the macroscopic growth behaviour and dynamics of individual water kefir grains throughout the fermentation process have not been investigated to date. Previous studies have primarily quantified changes in total grain mass at discrete time points under varying conditions [28,37,48,50,51]. While these studies demonstrated an increase in grain mass, weighing itself is not compatible with continuous monitoring. It requires either removing the grains from the fermentation batch, thereby interrupting the process, or setting up separate batches for each measurement time point. The method’s temporal resolution is limited by the inherent measurement interval, resulting in the omission of events occurring between weighing points.

Several studies have addressed aspects of grain formation, including proposed mechanisms of biofilm formation in milk kefir grains and the surface characteristics of biofilm-producing isolates derived from milk kefir [52–54]. For water kefir, the aggregation and network-forming behaviour of yeast cells in the presence of EPS-producing LAB was investigated as a contribution to the stepwise formation of granular structures [55]. These previous studies have expanded our knowledge at the micro- and batch scales. However, they have not addressed the macroscopic growth behaviour of individual grains.

Taken together, water kefir has become an increasingly popular fermented beverage worldwide, yet scientific knowledge remains comparatively limited beyond its fermentation technology and microbial community composition. Here, we studied the structural organisation, growth behaviour, and microbial community dynamics that jointly shape the propagation and biogeography of water kefir grains. By combining amplicon sequencing profiling with electron microscopy, we characterised the overall microbial composition and the internal architecture of the grains. Using a custom-built imaging system for time-resolved macroscopic documentation, we further analysed growth patterns in water kefir grains and liquid in parallel. To resolve the distribution and dynamics of microbial subpopulations, we complemented these approaches with a workflow integrating flow cytometry, fluorescence-activated cell sorting (FACS), and downstream amplicon sequencing. Taken together, these methods enabled the examination of grain structure, microbial organisation, and macroscopic growth behaviour across multiple scales, supporting the view that grain growth and propagation arise from the organisation of the multispecies aggregate rather than from its individual components alone.

## RESULTS

This study approaches water kefir from the perspective of its emergent multicellular functions, highlighting its potential as an accessible, but so far understudied, example of aggregative microbial multicellularity. We will focus on three interconnected characteristics of water kefir grains in a comprehensive and correlative manner, and to a resolution that has so far never been achieved:

i. The composition of our water kefir culture, both grains and liquid, was first analysed by conventional amplicon sequencing. These sample-specific analyses of ‘our’ water kefir community were complemented with a comparative *in silico* meta-analysis of data from 69 different samples of water kefir extracted from a recently published study [32]. This approach allowed the identification of predominant marker species commonly found in water kefir, as well as sample-specific microbial species. However, these results provide only a snapshot of the composition at the moment of sampling. To gain a quantitative, time-resolved picture of population/composition dynamics over the course of a fermentation cycle, we applied flow cytometry and fluorescence-activated cell sorting, followed by amplicon sequencing of the resulting gated subpopulations, to obtain a highly resolved, dynamic picture of the water kefir community.
ii. Water kefir growth has so far only been studied in bulk, e.g., by measuring the increase of biomass over time. Here, we developed a custom-built imaging system that enables continuous monitoring of individual water kefir grains throughout a 72-h fermentation cycle. These studies revealed an emergent multicellular function of kefir grain growth that was correlated with and can be explained by
iii. the ultrastructure of water kefir grains. Although some exemplary electron microscopy images from various regions of these macroscopic structures were available, an understanding of the water kefir grain architecture remained lacking. We combined stereomicroscopy with scanning electron microscopy (SEM) and transmission electron microscopy (TEM) to generate high-resolution micrographs of sections of whole water kefir grains. This structure of water kefir grains resembles microbial ‘planets’ with an uninhabited core of matrix components and cell debris, while living cells are only found in the outer shells and on the surface of the grains. This structure provides a biophysical explanation for the growth behaviour observed for individual water kefir grains, representing an emergent multicellular function of grain architecture and microbial growth.

### Integrated microbial profiling reveals distinct grain- and liquid-associated communities and sample-specific features of water kefir

DNA samples from water kefir grains and liquid were analysed by sequencing hypervariable regions of the bacterial 16S rRNA gene (Figure 1A) and the fungal ITS2 region (Figure 1B) to gain an overview of the microbial composition of the investigated culture. Some taxa could not be unambiguously resolved to a single species and are therefore reported using combined species names.

**Figure 1.**
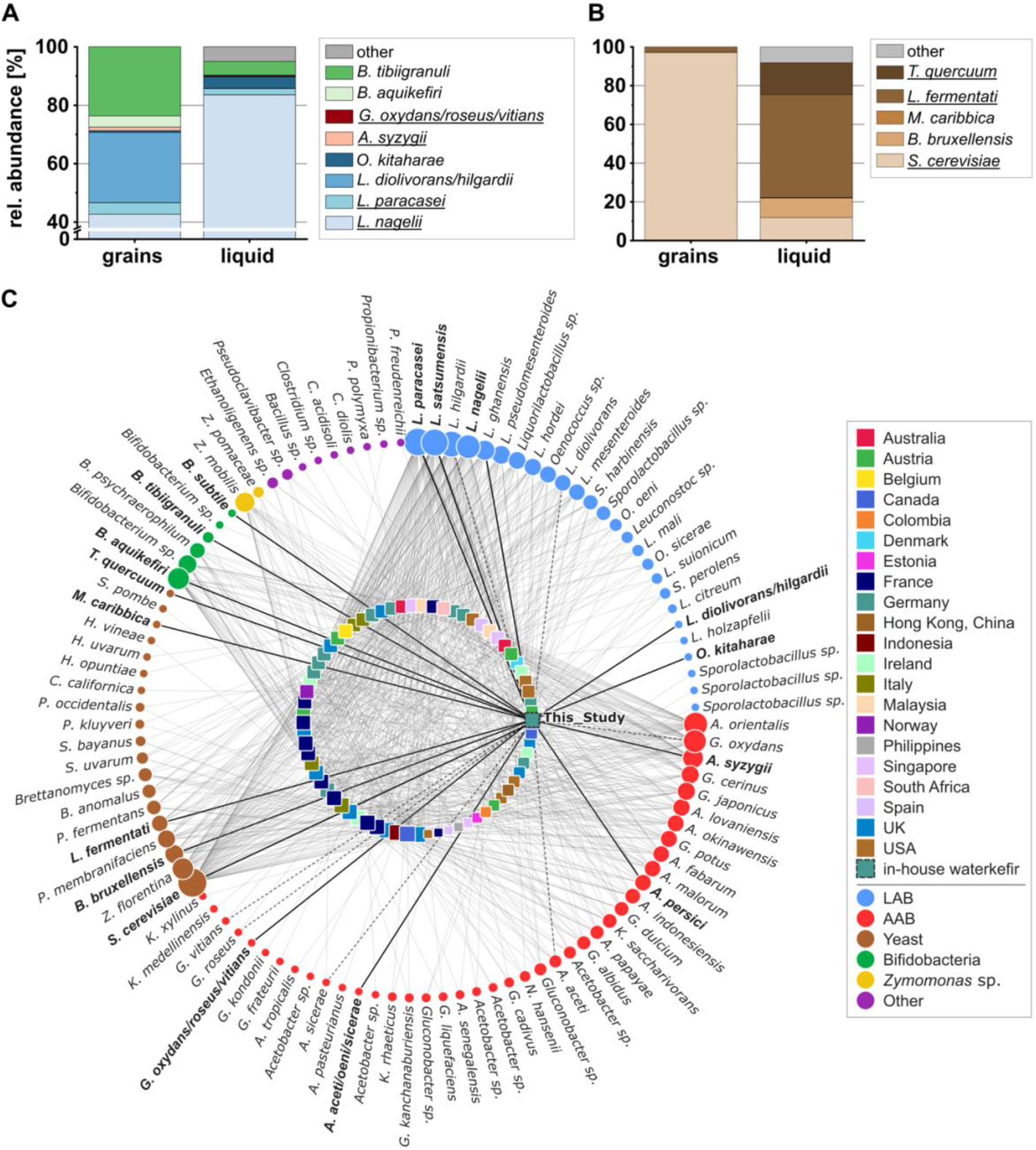
Microbial composition of the investigated water kefir culture and comparison with published water kefir communities. Relative abundances of (**A**) bacterial and (**B**) fungal taxa in kefir grains and in the liquid phase after 24 h of fermentation are shown based on OTU clustering. Combined species names indicate that the respective sequence could not be reliably resolved to a single species based on the amplified marker region. Underlined taxa indicate species that were successfully isolated in culture and added to our strain collection. (**C**) Radial bipartite network comparing the species profile of the water kefir culture analysed in this study with the 69 water kefir samples reported by [32]. Square nodes in the inner circle represent individual water kefir samples, including the culture analysed in this study; colours indicate country of origin, and node size reflects species richness. Circular nodes in the outer circle represent identified microbial species, with circle size indicating species prevalence across the complete dataset. Edges connect samples to the species detected in the respective culture. Edges connected to the culture analysed in this study are highlighted by increased line width. For ambiguously assigned taxa, dashed edges indicate links to the possible individual species.

After 24 h of fermentation, the grain and the liquid phase were dominated by *Liquorilactobacillus nagelii* (43% and 84%, respectively). The liquid additionally contained minor proportions of *Lacticaseibacillus paracasei* (2%), *Oenococcus kitaharae* (4%), and *Bifidobacterium aquikefiri* (5%), representing less abundant community members. The grains were more balanced, with higher proportions of *Lentilactobacillus diolivorans/hilgardii* and *B. aquikefiri* (both around 24%). AAB, such as *Acetobacter syzygii* and *Gluconobacter oxydans/roseus/vitians*, occurred only in trace amounts in both phases (<2%).

The yeast community was less diverse. *Saccharomyces cerevisiae* (97%) dominated the grains, whereas the liquid also contained *Lachancea fermentati* (57%), *Torulaspora quercuum* (17%), and *Meyerozyma caribbica* (0.3%), besides *S. cerevisiae* (12%). *Brettanomyces bruxellensis* was present in both phases but was more abundant in the liquid (0.5% and 10%, respectively).

In both bacterial and fungal analysis of the liquid, a small number of low-abundant sequences that were not typical for water kefir were identified and classified as ‘other’. These might be derived from the surfaces of the dried fruits, which served as a nitrogen source in our kefir cultivation medium. The detection of *T. quercuum* and *M. caribbica* may likewise indicate substrate-associated yeasts that became established at low abundance in the liquid phase under the cultivation conditions used here, rather than typical core members of the water kefir community. Their repeated detection in later FACS-based analyses further suggests that they were not merely sporadic contaminants but persisted as low-abundance members of the community.

In addition to this sequencing approach, several species present in the amplicon sequencing data were also isolated from grains and liquid, identified, and added to our collection of kefir isolates (Figure 1A and B; underlined taxa). Most importantly, isolates of *L. nagelii* and *Liquorilactobacillus satsumensis* produced large amounts of EPS on MRS agar supplemented with sucrose (Figure S3).

To place the microbial composition of our water kefir culture into a broader comparative context, we compared it *in silico* with the 69 water kefir samples recently described by [32]. For our culture, a consolidated species profile was generated by integrating amplicon sequencing, cultivation-based isolation, and FACS-based sorting, followed by sequencing. The resulting radial bipartite network visualises the distribution of identified microbial species across all samples, including the culture analysed in this study (Figure 1C). It shows that our culture harboured several microorganisms that were also frequently detected across published water kefir metagenomes, including representatives of the dominant LAB, AAB, and yeasts typically associated with water kefir fermentation (e.g., *L. paracasei, A. syzygii, S. cerevisiae*). In contrast, several taxa that occurred at low relative abundance in our sequencing data also showed low prevalence across the comparative dataset or were not detected in any other culture in the comparative dataset, including *O. kitaharae*, *Bifidobacterium subtile*, and *M. caribbica*. These species contributed to the sample-specific profile of our culture and underline that, beyond a shared core microbiome, individual water kefir communities can harbour distinct accessory members.

To further quantify sample-level similarity, we generated a water kefir similarity network in which individual samples, including our kefir culture and those from [32], were connected based on shared species composition (Figure S4). Network analysis revealed a highly connected structure, indicating that most water kefir samples shared at least a subset of microbial species. The culture analysed in this study showed the highest similarity to another German water kefir sample (WK024). The maximum conservative Jaccard similarity of our culture to any reference sample was *J_c_ = 0.33*, while the relaxed comparison, which accounts for ambiguously assigned taxa by considering overlap with possible species assignments, increased the maximum similarity to *J_r_ = 0.40*. Thus, even the closest match in the reference dataset showed only moderate similarity. This demonstrates that shared core taxa connect related samples, but that accessory or sample-specific taxa substantially contribute to the distinct microbial profile of our water kefir culture. Furthermore, WK024 remained the single most similar reference sample under both the conservative and relaxed Jaccard comparisons, and four of the five nearest samples were shared between the two metrics, indicating that the overall placement of our culture within the similarity network was largely robust to ambiguous taxonomic assignments.

### Fermentation dynamics and grain mass increase during bottle-scale water kefir fermentation

First, we aimed to document the growth behaviour and composition of water kefir grains using traditional approaches. We used a slightly modified growth setup in which the dried fruits were enclosed in paper tea bags and added to the fermentation bottles, allowing them to be easily removed before sampling. Throughout a seven-day fermentation, total soluble solids (TSS), pH, optical density at 600 nm (OD_600_), and grain wet mass were measured to monitor physicochemical changes and microbial growth (Figure 2A).

**Figure 2.**
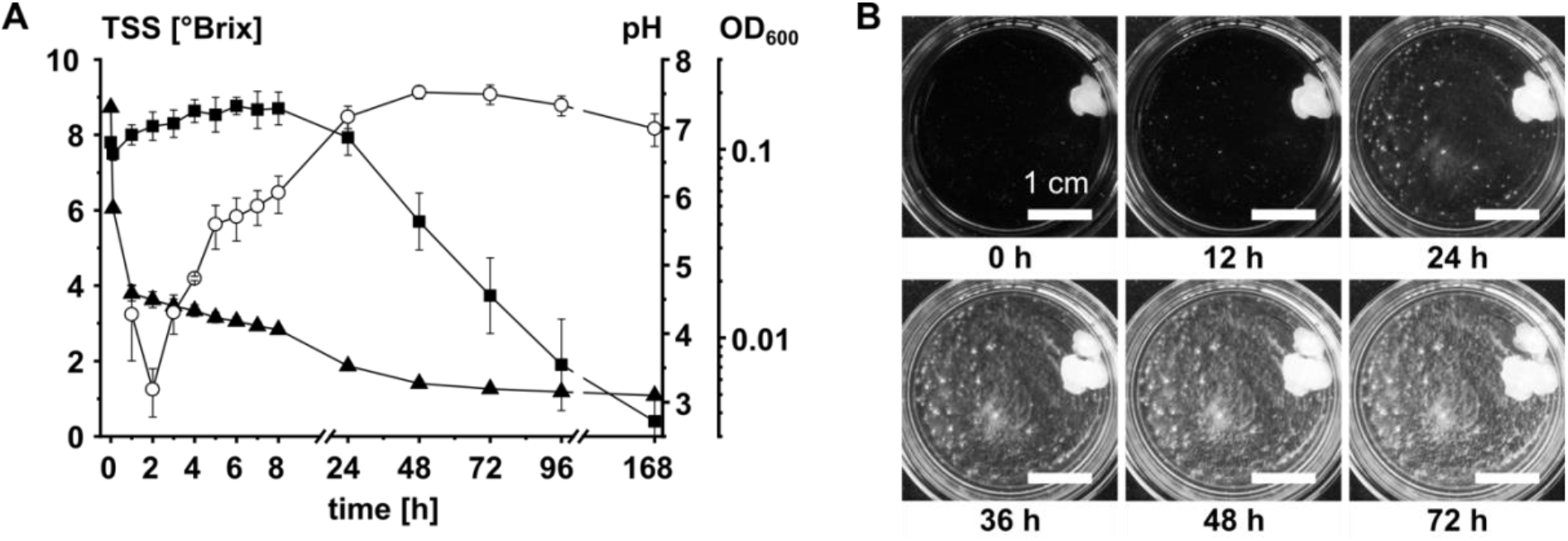
(**A**) Changes in fermentation process parameters, including TSS (■), pH (▲), and OD_600_ (○), in the water kefir liquid during a seven-day bottle-scale batch fermentation (*n* = 3). Error bars indicate standard deviation. (**B**) Representative images in 8-bit format, illustrating the macroscopic growth behaviour of a single water kefir grain in a miniaturised fermentation setup. Image acquisition was performed continuously over a 72-h observation period. A large-sized grain was inoculated in water kefir medium. Scale bars represent 1 cm.

The TSS showed a brief initial rise followed by a continuous decline, reaching nearly zero after 7 days. The pH dropped sharply after grain addition to the medium and stabilised around 3.2 within 48 h. Microbial growth in the liquid phase was monitored by measuring OD_600_ of the grain-free fermentation liquid. Initially, the value decreased during the first hour after grain addition, as small, flake-like conglomerates detached from them and gradually settled to the vessel bottom. OD_600_ then increased over the first 2 days before declining slightly again. The average grain mass increased from 51 g at inoculation to 64 g after 7 days. These results are in good agreement with comparable studies on the fermentation process parameters of water kefir [28,31,50].

Based on these results, the sampling times for the subsequent flow cytometry experiments were determined. Overall, the greatest changes in the measured process parameters occurred within the first 24 h of fermentation, supporting the dense sampling intervals during this period for subsequent flow cytometric analysis. Thereafter, pH and OD_600_ changed only slightly, whereas TSS continued to decrease, providing the basis for daily sampling until day 4. The additional sampling point on day 7 was included to assess whether prolonged exposure to acidic and increasingly nutrient-limited conditions led to further changes in the microbial community.

### Flow cytometry reveals stable grain communities and dynamic changes in the liquid throughout fermentation

To assess structural dynamics of the microbial community during water kefir fermentation, time-resolved samples from both grain and liquid fractions were analysed by DAPI-stained flow cytometry (for process control parameters, see Figure S5 and Table S2). The resulting 2D plots show DAPI fluorescence, which reflects DNA content, plotted against forward scatter (FSC), a measure of cell size (Figure 3A, Figure S6). A total of 31 cytometric gates (G1–G31) were defined and applied across all samples. In the case of studying water kefir, gates in the high-FSC/high-fluorescence region, typically enriched in larger, yeast-sized cells but also small bacterial aggregates, accounted for six gates. In contrast, the remaining 25 gates, characterised by lower FSC and fluorescence, corresponded to smaller, bacterial-sized cells. These groups are based on cytometric properties and do not imply taxonomic exclusivity. Figure 3A illustrates the applied gate template and representative cytometric patterns for grain and liquid samples from one biological replicate during one fermentation cycle. While the grain samples showed largely similar event distributions over time, the liquid samples displayed more pronounced changes in event density across several gates. Events in the high-FSC/high-fluorescence region tended to decrease over time, whereas events in the low-FSC/low-fluorescence region increased during the first day before declining again.

**Figure 3.**
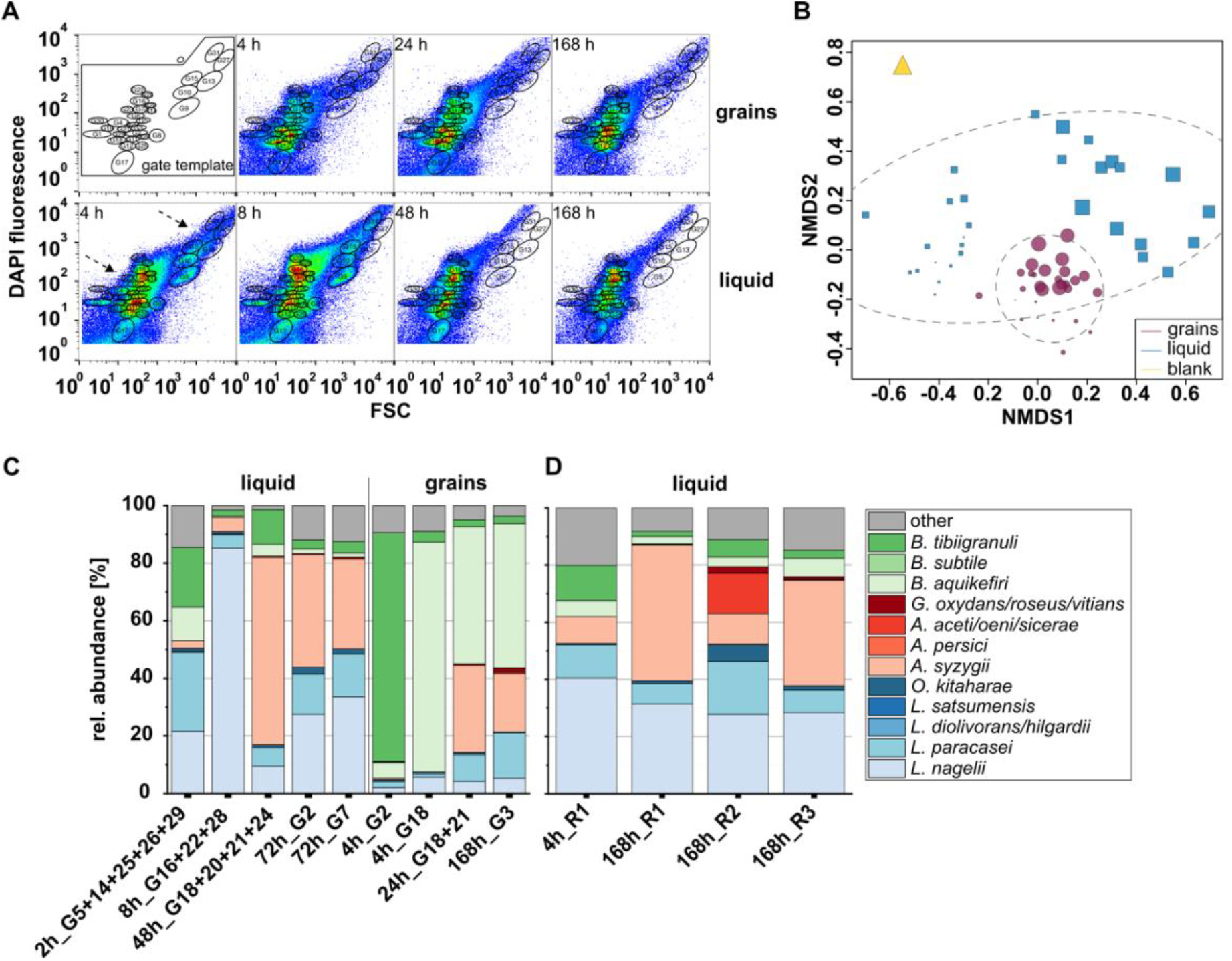
Cytometric gating, community structure, and taxonomic profiling during water kefir fermentation. (**A**) The gate template, consisting of 200,000 cells and excluding instrumental noise and beads, defined 31 unified gates across all samples. Representative flow cytometry 2D plots of DAPI fluorescence at 460±25 nm (355 nm) versus forward scatter (FSC) at 488 nm are shown for grain and liquid samples at selected time points, as indicated in each plot. Dashed arrows point at the low and high FSC/DAPI regions in the liquid samples in which pronounced changes in event density were observed over time. (**B**) NMDS plot based on cytometric gate composition, showing grains (●), liquid (■), and substrate blank (▲). Larger symbol sizes correspond to later sampling time points within the 168-h fermentation. (**C**) Taxonomic profiles of selected subcommunities that showed high or consistently abundant representation in the liquid or grain fractions over time. Sample labels indicate the sampling time point during fermentation, followed by the sorted single gate or gate combination used for subsequent amplicon sequencing. Relative abundances of pooled ASVs are shown at the species level. Combined species names indicate that the respective sequence could not be reliably resolved to a single species based on the amplified marker region. (**D**) Taxonomic profiles of 16S amplicon sequencing results from all low FSC/low DAPI gates in the liquid fraction at 4 h (replicate R1) and at 168 h (replicates R1–3), illustrating temporal shifts and endpoint variation among biological replicates of bulk fermentations.

Community dynamics were further assessed using non-metric multidimensional scaling (NMDS; Figure 3B) to track structural differences between samples from liquids and grains. Sample information was obtained by cytometric measurement of the respective samples and the evaluation of the cytometric 2D plots (Figure S6) using the tool flowCyBar [56]. All grain samples exhibited high structural stability throughout fermentation. The NMDS distribution showed tight clustering, with only a slight shift between early (0–8 h) and later (24–168 h) time points. Similarly, the flowCyBar heatmap of the grain-associated gates (Figure S7A) showed that their relative abundances underwent negligible changes, with a trajectory towards gates G3, G5 and G20 by the end of the experiment (168 h). Some gates (e.g., G2 and G18) exhibited high relative abundances (around 11% and 9%, respectively) throughout the entire fermentation process, and only a few (e.g., G3 and G21) showed strong time-dependent variation. Correlation analysis (Figure S7C) identified a limited number of strong interactions, primarily positive, that occurred between gates (e.g., G18 and G21; G14, G29, and G31) rather than with abiotic parameters. Only one gate (G3) correlated significantly negatively with the pH as an environmental factor.

In contrast, liquid samples underwent pronounced temporal dynamics. The NMDS plot showed a wide spatial distribution, with clear separation between early (up to 24 h) and late (48–168 h) samples. Notably, the three biological replicates at 168 h exhibited substantial internal variation. The heatmap (Figure S7B) verified this pattern, with fewer consistently dominant gates, e.g., G2 and G7 (around 11% and 10%, respectively), and greater temporal shifts in relative abundances, e.g., G5, G10, G13, G14, G15, G25, G26, G27, G29, and G31. In the liquid, distinct gates showed fermentation phase-specific abundance maxima. Gates G5, G14, G25, G26, and G29 peaked during the early phase at 2 h, whereas G16, G22, and G28 reached their highest abundance at the intermediate 8 h time point. Later in fermentation, at 48 h, maximum relative abundances shifted to G18, G20, G21, and G24. Correlation analysis (Figure S7D) revealed numerous strong interactions, both positive (e.g., G16, G22, and G28) and negative (e.g., G18 and G31). Several gates showed positive and negative correlations with the parameters pH, OD_600_, and TSS (e.g., G21, G25, and G26). The strongest effects of these environmental factors were observed during the early fermentation phase, characterised by a high TSS level and a substantial pH drop.

For deeper community profiling, selected gates were sorted, either individually or as predefined gate combinations, and analysed by 16S or ITS amplicon sequencing. A substrate control sample consisting of water, dried figs, and sugar was clearly distinct from the fermentation samples (Figure 3B). The amplicon sequencing analysis revealed a diverse microbial composition, including bacteria and fungi typically associated with plants or soil. This highlighted the microbial load introduced by substrate components, particularly unpreserved dried fruits (Figure S8A and B). Bacterial reads were dominated by *Rhizobiaceae* and *Xanthobacteraceae*, with high overall diversity but uneven taxonomic distribution. Notably, the fungal fraction was dominated by an amplicon sequence variant (ASV) of *S. cerevisiae* that differed from the strain found in the kefir samples.

To assess how cytometrically enriched regions relate to microbial taxonomy, all gates from the low-FSC/low-DAPI (smaller, low-fluorescent cells) and high-FSC/high-DAPI (larger, high-fluorescent cells) areas were pooled and sorted for 16S and ITS sequencing. Both bacterial and fungal taxa were detected in all regions (Figure S8C and D), although event density was substantially higher in the low-FSC/low-DAPI fraction.

Fungal subcommunity composition was further characterised by sorting six individual gates from the high-FSC/high-DAPI region of grain or liquid samples (Figure S8E). In grains, gates G13 and G27 were dominated by *S. cerevisiae*, with minor *B. bruxellensis* contributions and non-kefir reads in G13, likely originating from residual substrate or nonspecific cell associations in early samples. Their relative abundances peaked during early fermentation. In liquid, gates G9, G10, G15, and G31, also sorted from early time points, showed greater diversity, dominated by *B. bruxellensis* and *M. caribbica*, along with lower levels of *S. cerevisiae* and *L. fermentati*. To capture broader patterns at 8 h, abundant gates were combined by fraction (Figure S8F). Grain gates remained dominated by *S. cerevisiae*, while liquid gates revealed a more even distribution of the four dominant yeasts: *S. cerevisiae*, *B. bruxellensis*, *L. fermentati*, and *M. caribbica*, with minimal non-kefir or unassigned taxa.

Bacterial subcommunity dynamics differed markedly between liquid and grain fractions (Figure 3C). In the liquid, early-stage samples were dominated by LAB, especially *L. nagelii* and *L. paracasei*, while *Bifidobacterium* spp. were present in smaller proportions, and AAB were initially rare. Over time, *A. syzygii* became dominant, although *Lactobacillus* spp. and *Bifidobacterium* spp. remained detectable. Gates G2 and G7, which showed the highest average relative abundances, displayed balanced communities composed of 40% LAB, 40% AAB, 10% *Bifidobacterium* spp. And *O. kitaharae*, with the remainder consisting of non-kefir species likely introduced via the substrate. In the grain fraction, gates G2 and G18 were consistently abundant and dominated by *Bifidobacterium tibiigranuli* in G2 and *B. aquikefiri* in G18. A combined sort of G18 and G21 at 24 h revealed a mixed community of *B. aquikefiri*, *A. syzygii*, and *Lactobacillus* spp.. Gate G3, which showed the highest relative abundances in grains at 96 h and 168 h, was also sorted and exhibited a similar profile, with additional *G. oxydans/roseus/vitians* detected at 168 h.

To assess the functional consequences of bacterial community shifts, low-DAPI/low-FSC gates were sorted at 4 h from replicate 1 and at 168 h from all three biological replicates (Figure 3D). At 4 h, the community was composed of 50% *Lactobacillus* spp., with the remainder split between *Bifidobacterium* spp. And *A. syzygii*. A considerable fraction of non-kefir-associated sequences was also detectable. By 168 h, all replicates showed a clear transition: *Lactobacillus* spp. abundance decreased, while *A. syzygii* became dominant. Replicate 2 additionally contained higher levels of other AAB, including *Acetobacter aceti/oeni/sicerae*, and *G. oxydans/roseus/vitians.* Non-kefir taxa declined, and *Bifidobacterium spp.* were reduced but still present, while *O. kitaharae* increased across all replicates. The compositional differences at the endpoint appear to result from shifts in species proportions within functional groups, rather than from complete taxonomic turnover.

### Time-lapse imaging uncovers the macroscopic growth behaviour of water kefir

The well-documented increase in biomass is accompanied by an increase in kefir grain volume, which also correlates with a higher number of water kefir grains. But the mechanism underlying this increase is poorly understood. Towards this end, we developed and optimised a custom-made imaging system that enables monitoring kefir grain development in a miniaturised cultivation setup throughout the complete growth cycle under conditions that mimic standard fermentation (see methods and supplemental material for detailed information and protocols).

Over the 72-hour observation period, continuous imaging yielded more than 860 frames per experiment. The resulting image sequences capture the growth behaviour and structural changes of the kefir grains and the surrounding liquid. Access to time-lapse videos is provided in the supplementary material.

A series of experiments was performed in which the inoculation volume was regulated by either the initial grain size (Figure 2B, Figure S10, Videos 1–4) or the number of grains per sample vessel (Figure S11, Videos 5–7). Throughout all set-ups, the grains continuously increased in size as fermentation progressed. However, this growth did not occur uniformly in all dimensions. Instead, slight surface expansion asymmetries often resulted in irregular shapes. The indentations and cavities that accumulated in the growing grains eventually caused them to break apart (Figure 2B, Video 1 and Video 2). This process appears to be the key mechanism of grain multiplication. No *de novo* formation of macroscopically detectable grains was observed in the medium throughout any of the experimental series. The observed ‘movement’, particularly in large and extra-large kefir grains, was caused by gas bubble formation as a result of fermentation activity (Figure S10, Videos 2–4).

In addition to grain growth, turbidity of the surrounding liquid increased, presumably due to cells originating from the grains and subsequently multiplying in the liquid phase. Light microscopy revealed that these were primarily bacterial cells, with yeast cells also present (Figure S12). In the absence of agitation, suspended cells gradually settled to the bottom of the vessels, forming a biofilm. Biofilm formation tended to decrease with increasing inoculum, both with increasing size of individual grains and increasing number of grains. Additionally, white flakes up to 1 mm in size were consistently observed in the liquid shortly after addition of the medium. Similar structures were also observed in bulk fermentations. These cell conglomerates consisted of bacterial and yeast cells (Figure 5F) and showed slight growth during the experiments but did not develop into macroscopically detectable grains within the observation period. Whether these structures represent transient cell aggregates or potential early stages of *de novo* grain formation could not be determined from the present experiments.

### Grain and liquid phase follow distinct growth trajectories

To gain more quantitative insights into the growth process, white pixel data were extracted from the time-lapse images and analysed. All image frames were processed using a predefined set of three regions of interest (ROIs): a total ROI covering the entire sample area, a grain ROI capturing the area associated with the grains, and a liquid ROI covering the area surrounding the grains (Figure S2). These ROIs were applied to all time points, thereby ensuring spatial comparability and enabling the separate quantification of grain- and liquid-associated growth signals.

Tracking the number of white pixels over 72 h in the total ROI revealed typical sigmoidal microbial growth curves comprising lag, exponential, and stationary phases (Figure 4A, Figure S13, Figure S14A and D). Initial pixel values increased with the number or size of inoculated grains, leading to different absolute signal levels at the start of the experiments. By contrast, endpoint values converged within a similar range, suggesting that the total ROI approached saturation within the 72-h observation period, independent of the initial grain number.

**Figure 4.**
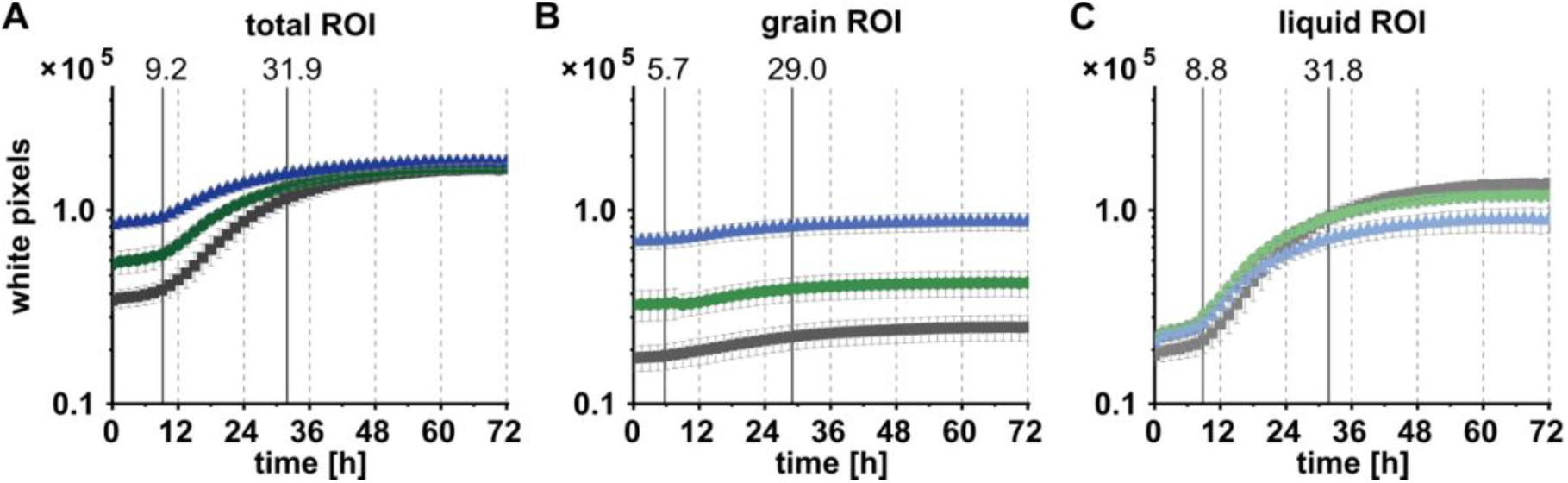
Quantitative analysis of image-derived growth dynamics at different initial grain numbers. (**A–C**) Representative growth curves showing the increase in white pixel counts over time for samples inoculated with 2 (■), 4 (●), and 8 (▲) medium-sized grains. Curves represent the mean values from three biological replicates per condition in a single 72-h experiment. Images were analysed using the (**A**) total ROI, (**B**) grain ROI, and (**C**) liquid ROI. Vertical, solid lines indicate the median time points at which 5% and 70% of the absolute increase in white pixels was reached across all samples in one experiment. Error bars indicate standard deviation.

The sigmoidal pattern observed in the total ROI was mainly reflected by the liquid ROI, which showed the greatest dynamic changes over time (Figure 4C, Figure S14C and F). In contrast, the grain ROI exhibited only a slight, almost linear increase (Figure 4B, Figure S14B and E). This weaker increase likely reflects the limited ability of two-dimensional image-based measurements to capture the volume expansion of three-dimensional grains within a predefined ROI. However, the method validation showed good agreement between image-derived two-dimensional signals and three-dimensional grain growth, supporting the use of white pixel counts as a suitable proxy for growth-related changes (see supplemental results for method verification).

In general, liquid-associated growth followed a similar pattern to total growth but appeared delayed compared with the signal increase in the grain ROI. Thus, the liquid phase largely shaped the temporal course of the total growth curves, whereas the number of inoculated grains mainly determined the initial signal level. A higher number of starter grains increased the initial white-pixel signal. It accelerated the early rise in the liquid-associated signal but also led to earlier saturation of the total ROI, thereby constraining further signal increase toward the end of the observation period.

The empirically defined thresholds of 5% and 70% of the total increase in white pixels corresponded, respectively, to the transitions from the lag to the exponential and from the exponential to the stationary growth phase in the total ROI. For consistent comparison, the same thresholds were also applied to the grain and liquid ROIs and are indicated by vertical lines Figure 4. The corresponding median time points were approximately 9 and 32 h for the total and liquid ROIs, whereas both thresholds were reached earlier in the grain ROI, at approximately 6 and 29 h, respectively. The corresponding threshold times, together with other growth parameters derived from the pixel data, are quantitatively compared in Figure S15 in the supplemental material.

### Water kefir grains exhibit a gradual internal transition from cell-rich to matrix-dense regions

To gain a comprehensive understanding of water grain architecture throughout all size scales, we combined different microscopic techniques to unravel the macro- and microstructure of kefir grain surfaces, as well as fully resolve the ultrastructure of complete cross-sections of individual water kefir grains (Figure 5). This combined approach yielded fundamental and highly surprising insights into grain architecture, which also provides a possible explanation for the growth behaviour described above (Figure 4).

**Figure 5.**
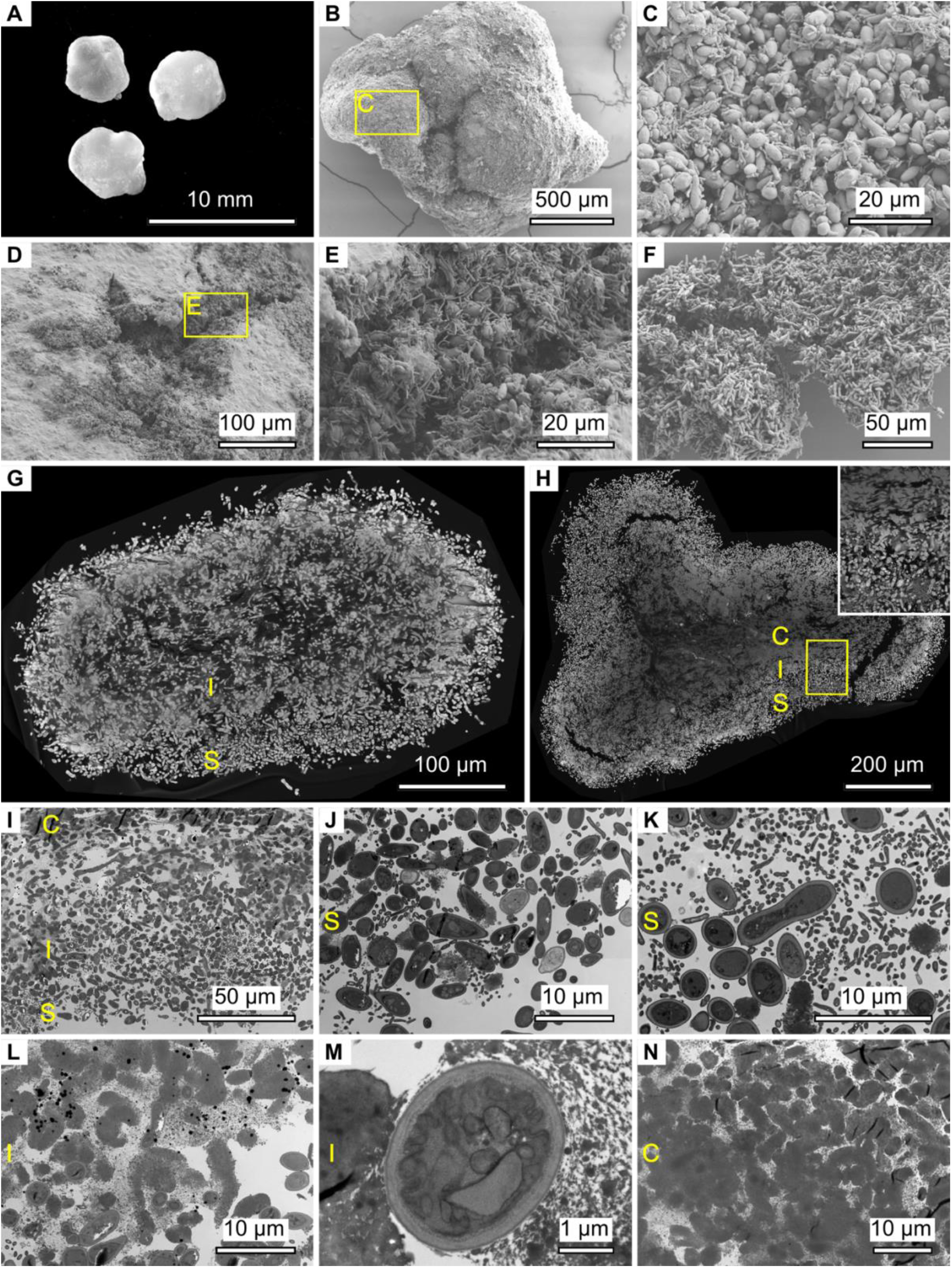
Macro- and micro-morphology of water kefir grains. (**A**) Macroscopic view of medium-sized water kefir grains. (**B–E**) Scanning electron micrographs showing the surface and an indentation of a grain: (**B**) overview of an intact grain, (**C**) close-up of the outer surface covered with bacteria and yeasts, (**D**) indentation inside a cut-open grain, (**E**) surface of the indentation densely populated with microbial cells. (**F**) A conglomerate of detached yeast and bacterial cells from the grain surface. (**G, H**) Backscatter images obtained using scanning electron microscopy of cross-sections through an embedded water kefir grain. Identified structural regions are labelled: S - surface, I - interphase, and C - core: (**G**) Section close to the surface, and (**H**) section through the central region of the grain. (**I–N**) Transmission electron micrographs of a cross-section through a water kefir grain: (**I**) overview showing the three structural region (S, I, C), (**J**) close-up of a yeast-rich subarea within the surface region (S), (**K**) bacteria-rich subarea within the surface region (S), (**L**) interphase region (I) with emerging extracellular matrix and intact cells, (**M**) comparison of an intact cell (centre) with a cell remnant (left) and extracellular EPS matrix (right) in the interphase region (I), and (**N**) core region (C) containing EPS matrix and enclosed cell remnants.

The macro- and micromorphological characteristics of water kefir grains were analysed using stereomicroscopy (Figure 5A), SEM (Figure 5B–H), and TEM (Figure 5I–N). On a macroscopic level, grains appear largely spherical with irregular surfaces. They are typically white and appear more translucent in thinner areas. Their size ranges from 1-3 mm (smallest, Figure S16A) to 10 mm and above (largest Figure S16B), with the vast majority falling between 4-6 mm (Figure 5A).

SEM analysis showed a dense microbial surface layer dominated by yeasts, with bacteria interspersed and occasionally surrounded by EPS, but mostly exposed to the environment (Figure 5B and C). Indentations extending into the inner matrix were colonised mainly by long, rod-shaped bacteria with fewer yeasts than on the surface (Figure 5D and E). In addition to compact grains, free-floating white flakes were observed in bulk fermentations in the liquid phase. Similar structures were also documented in grain growth studies in the miniaturised setup (Figure 2B, Figure S10, Figure S11). As revealed by SEM analysis, these microbial conglomerates consisted of loosely aggregated yeast and bacterial cells but lacked both a substantial EPS matrix and a dense core (Figure 5F). Notably, yeast cells in the flakes appeared more elongated than the oval forms typically found on the grain surface.

Representative sections through the peripheral (Figure 5G) and central (Figure 5H) grain regions were generated by stitching together multiple individual negative-stained SEM micrographs of embedded grain sections. This approach allowed the ultrastructure of water kefir grains to be unravelled for the first time as an emergent multicellular entity. It revealed three morphologically distinct regions: surface, interphase, and core, with gradual transitions. To examine the structure of the individual regions in greater detail, ultrathin sections were prepared and examined using TEM (Figure 5I-K). As observed in SEM imaging of intact grains, the surface consisted of dense microbial layers, including yeast- and bacteria-dominated regions (Figure 5J and K). The layers varied in thickness and sometimes extended inward, consistent with the previously described indentation. Most cells had intact cell walls, indicating they were viable before fixation. The presence of fibrous material attributed to bacterial EPS in this surface region was rare, confirming that it is primarily cell-rich rather than matrix-dominated.

In contrast, the underlying interphase region consisted of network-like EPS, and additional structures resembling yeast in shape and size (Figure 5L). Unlike viable cells on the surface, these remnants lacked intact cell walls and internal structures. Some intact cells were still present, distinguishable by their defined shape and intact envelope, but they became increasingly embedded in the matrix (Figure 5M). The core represented the final structural shift from the cell-rich periphery to a purely matrix-dominated interior. It consisted solely of EPS and cellular debris, with no intact and potentially viable cells remaining (Figure 5N). Remnants were progressively deformed and compressed into dense aggregates. A comparison of the two cross-sections (Figure 5G and H) showed that the outer section consisted predominantly of the surface and interphase regions, whereas the inner section exhibited a distinct core region (Figure 5H), characterised by a greater loss of defined cellular structures.

Taken together, the ultrastructure revealed that water kefir grains represent a rather unique type of microbial ‘planet’: an emergent multicellular structure formed by yeast and bacteria, inhabited only in the outer shell, with cell debris and a compacted matrix forming a lifeless inner core. This unique structure may explain the unusual multicellular way of growth observed in our images (Figure 2B, Figure S10, Figure S11) and time-lapse videos (see supplemental material).

## DISCUSSION

Water kefir grains thrive in a dynamic environment that requires microorganisms to adapt continuously. This study focused on growth as a specific emergent multicellular function of the structured multispecies community within water kefir grains. They represent a prime example of aggregative multicellularity, forming biofilm-like grains in which bacteria and yeasts coexist by shaping their own habitat. Several characteristics define this system: the grains exhibit a specific metabolic profile with high stability [57], they are more resilient than planktonic cultures [58], and their growth depends on the enlargement and fragmentation of existing grains, as shown in this study. Once established, there is no need for the multispecies community in the grains to return to early successional stages of community assembly [58]. While previous studies primarily focused on the community composition or on fermentation performance across varying substrates, a comprehensive, correlated analysis of structure, microbial dynamics, and growth behaviour has been lacking. This study closes this knowledge gap by combining a highly resolved structural characterisation with a quantitative time-resolved description of growth and multiplication phenomena at the single-grain level. The resulting comprehensive data allow the macroscopic behaviour of multicellular growth to be described as a function of both microbial composition and the microscopic organisation and architecture of water kefir grains (Figure 6).

**Figure 6.**
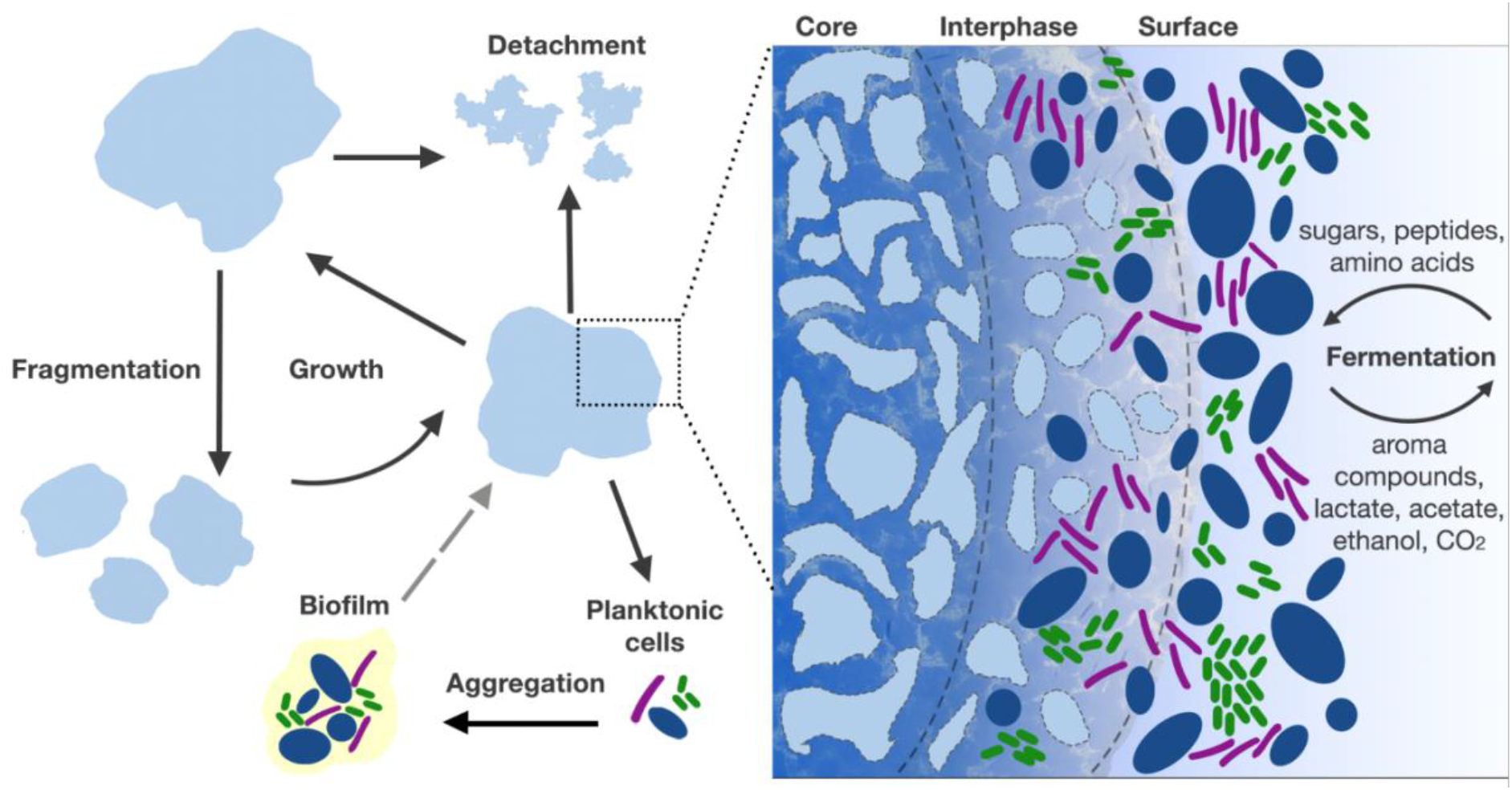
Schematic representation of the structural characteristics of water kefir grains and the associated growth phenomena during fermentation. The dashed arrow indicates a hypothetical, literature-based pathway for grain formation from biofilm structures [53]. Other potential routes could not be excluded within the observation period of this study.

The new imaging approach enabled the visualisation and quantitative analysis of water kefir growth from two-dimensional white-pixel data. This method reliably captures biologically meaningful growth dynamics, even within the complex, multispecies community. To our knowledge, this is the first method that tracks multiple macroscopic growth processes in both kefir grains and the surrounding liquid. Defining distinct ROIs for each phase provides additional insights into the interdependent dynamic processes that conventional measures, such as wet weight or optical density, cannot capture. Despite the potential for bias in samples containing extra-large or fragmenting grains, this method is a powerful, non-invasive tool for monitoring macroscopic microbial growth phenomena at high temporal resolution, providing direct visual evidence.

While earlier studies proposed a heterogeneous distribution of cells and EPS [47,49,59], subsequent publications often generalised this to describe kefir grains as EPS-based structures with microbial cells embedded in the matrix. By contrast, our findings, which are based on highly resolved cross-sections, revealed that kefir grains have a tripartite structure, with a gradual transition from a cell-dominated surface to a matrix-rich core. Grain growth itself presumably occurs in the surface region, where intact microbial cells are present and able to proliferate. After a brief lag phase, the macroscopic expansion was largely complete after 30 h in the miniaturised fermentation setup. Growth appears slower and less dynamic than that of planktonic cells in the liquid, suggesting gradual grain enlargement rather than unrestricted exponential growth. This difference may partly arise from the spatially constrained nature of grain-associated growth, in which cell proliferation and EPS matrix production occur within an existing three-dimensional structure, whereas planktonic cells can proliferate throughout the surrounding liquid. In addition, local gradients in nutrients, oxygen, and metabolic products within the grain may further constrain microbial activity. In the interphase, EPS accumulate and form network-like layers, potentially restricting nutrient availability and the space available for cell proliferation. Progressive nutrient limitation and eventual starvation may contribute to cell death towards the grain interior, consistent with the accumulation of dense matrix and cell debris observed in the core.

Envelope-free cellular remnants embedded in EPS have also been described for milk kefir grains [60]. Later, prolonged cultivation was shown to promote the accumulation of extracellular substances that enclosed cells and was accompanied by cell death, while nutrients released from dead cells could subsequently support the growth of neighbouring microorganisms [36]. Decades of studying microbial biofilms have established that the extracellular matrix not only consists of EPS but also contains significant amounts of additional components, such as amyloid proteins, extracellular DNA, and lipids, that result from cell lysis within the core matrix [61,62]. The observation reported herein, as well as the older reports in milk kefir, are therefore in good agreement with the vast body of knowledge on heterogeneity and structure of the biofilm matrix.

In the context of kefir grains, excessive growth would likely disrupt the matrix and weaken grain stability. But since the non-living core, with its firm gel-like texture, remains static, microbial division in the outer shell will generate increasing shear forces within the core, ultimately promoting fragmentation. This process, which we have documented at the single-grain level for the first time, presumably explains why the grains in normal bulk fermentations are predominantly of an average medium size, while very large grains are found only rarely.

The relationship between grain structure and growth may be understood as a feedback process rather than as a one-directional effect. Spatial organisation in microbial communities can emerge through feedback loops in which microbial activity modifies the local environment, which in turn affects subsequent growth and organisation [63]. In water kefir grains, microbial proliferation and EPS production generate a distinct architecture with a cell-rich surface and an increasingly compact interior. Once established, this architecture may favour continued growth at the grain periphery, while constraining expansion within deeper regions. Continued peripheral growth, in turn, further modifies grain architecture. Our findings are therefore consistent with a growth– architecture feedback in which microbial activity shapes the grain structure, while the resulting structure directs its further development.

The microbial kefir grain community displayed notable structural and compositional stability throughout the fermentation process, in line with the limited dynamics of the grain growth curves. Throughout growth, it was dominated by *S. cerevisiae* and *Bifidobacterium* spp., with only minor fluctuations over time. In addition to these core members, the grains are inhabited by other functional groups, including LAB and AAB, that spread out into the liquid phase after inoculation. The biofilm structure of the grains likely plays a key role in their structural and compositional stability. It provides physical protection against environmental stressors and offers a spatial scaffold that anchors the microbial kefir community to a stable, self-built, gel-like core.

In contrast, the fermentation liquid exhibited more pronounced dynamics in both growth behaviour and community composition, including a gradual shift from LAB to AAB. This succession likely reflects the changing environmental conditions during fermentation, including substrate depletion, acidification, and the accumulation of fermentation products, with ethanol produced by yeasts providing an additional substrate for AAB. The grain-associated community also changed over time, but these shifts were more subtle than those observed in the liquid fraction. The release of cells from the grain surface and their subsequent proliferation enabled colonisation of the liquid phase, leading to increasing turbidity and biofilm formation at the vessel bottom. Our findings complement previous observations of distinct dynamics in the grain and liquid fractions during early fermentation stages [30]. By extending the analysis to 168 h and combining flow cytometry with targeted amplicon sequencing of sorted subcommunities, we obtained a more resolved view of longer-term microbial dynamics.

So far, the *de novo* formation of kefir grains has primarily been associated with biofilm formation [36,52–55]. Since our experiments were performed with already established grains, our imaging data cannot directly address this hypothesis. However, they show that, under the conditions tested here, the multiplication of existing grains was not driven by *de novo* biofilm-derived grain formation in the liquid phase, but mainly by fragmentation of grains beyond a critical size or due to mechanical forces (Figure 6). Most likely, this is also the only relevant mechanism of water kefir growth in the typical fermentation setting, particularly, since washing the grains prior to the next round of kefir production and the use of clean, biofilm-free vessels is the standard procedure [33].

We also observed the presence and growth of macroscopically visible cell conglomerates in the liquid phase. Similar flake-like structures were found in batch fermentations, suggesting that they are not artefacts of the miniaturised setup. Comparable non-surface-attached microbial aggregates have been described as an organisational form between free-living planktonic cells and surface-attached biofilms, often involving the association of bacterial cells with EPS-rich particles [64]. The water kefir conglomerates differed from many of these examples by comprising both bacteria and yeasts and containing little apparent matrix material. Moreover, their abundance was strongly reduced after washing the grain (data not shown), suggesting that they originate, at least in parts, from material released from the grain surface rather than having formed *de novo* in the liquid phase. To our knowledge, such structures have not previously been described in water kefir. Although these conglomerates did not develop into compact grains under the tested conditions, they may nevertheless represent an additional growth form and potentially provide precursor structures for *de novo* grain formation under specific environmental conditions.

Taken together, our findings highlight a complex relationship between the grain and the liquid phase, in which distinct yet interconnected microbial communities shape the fermentation system as a whole. The grains provide a structurally stable biofilm-matrix that supports compositional continuity, growth, and resilience. They also function as a microbial reservoir from which planktonic cells are released into the liquid phase. The functional coupling of structural robustness and adaptive flexibility is a defining feature of the water kefir ecosystem.

Studying microbes as multicellular entities is the result of a relatively recent change in perception, supported by methodological advances that now allow macroscopic microbial structures to be resolved down to the single-cell level [65]. By combining continuous observation of individual grains with microscopic analysis of their cellular organisation, this study linked growth and propagation at the whole-grain level to spatial differences in cellular and matrix organisation within the grains.

Here, we propose water kefir grains as a novel, highly accessible model system for studying aggregative microbial multicellularity. Natural kefir grains are complex, submerged microbial biofilms that are not associated with a surface. These globular biofilms are composed of multiple species from overlapping functional groups that compete with, but also synergistically co-dependent on one another. Water kefir grains are easy to obtain and simple to cultivate. Moreover, they consist of microorganisms that may become accessible to genetic manipulation in the future, providing an opportunity to unravel the molecular mechanisms underlying emergent multicellular functions, such as the grain-specific mode of growth and division described here. With this study, we hope to provide a first step towards a holistic perspective of a promising new model for a multicellular microbial system.

## MATERIALS AND METHODS

### Water kefir cultivation

The water kefir grains were obtained from a private household (Dresden, Germany). Fermentation was carried out following a standard procedure: 64 g refined sucrose (8% wt/vol) were dissolved in 800 mL tap water in a screw-cap glass bottle, and 50 g kefir grains (6% wt/vol) were added. Dried figs without added preservatives (dmBio, Karlsruhe, Germany) served as the standard fruit; 32 g (4% wt/vol) were cut into eighths, placed in a tea bag, and suspended in the bottle. The tea bag ensured the complete removal of fig pieces after fermentation and was fixed to the bottle neck with a slightly opened screw cap to allow gas exchange. Fermentation was performed at room temperature (18–21 °C) and stopped every 2–3 days by discarding the tea bag and liquid. The grains were then sieved, rinsed with running tap water, and reused under identical conditions.

### Fermentation process parameters

The parameters pH, TSS, and OD_600_ were monitored during standard fermentations performed in triplicate as described above. On the first day, samples were taken hourly between 0–8 h, followed by daily sampling after 24, 48, 72, and 96 h (1–4 d) and again after 168 h (7 d). At each time point, 10 mL were collected; 1 mL was used to measure OD_600_, and the remainder was sterile-filtered (0.2 µm) for pH and soluble solids determination and used as the blank for OD_600_. pH was measured with a glass electrode (WTW, Weilheim, Germany), OD_600_ with an Ultrospec 2100 pro UV/Vis spectrophotometer (Amersham Bioscience, Amersham, United Kingdom), and TSS with a DMA 35 Basic Handheld Density Meter (Anton Paar, Ostfildern-Scharnhausen, Germany). The final grain wet mass was determined after 7 days by sieving, draining for 5 min, and weighing.

### Microbial community analysis

#### DNA extraction from water kefir grains and fermented liquid

A culture-independent species analysis was conducted to assess bacterial and yeast diversity. Genomic DNA (gDNA) was extracted from grains and liquid samples after 24 h of fermentation. Grains were rinsed thoroughly with tap water, drained for 5 min, and 5 g were freeze-dried. The dried material was ground with a mortar and pestle to a fine white powder, of which 50 mg were suspended in 1 mL phosphate buffer (50 mM, pH 6.5). Enzymatic hydrolysis was performed with 10 µL endo-dextranase (1:10 dilution in phosphate buffer; from *Chaetomium erraticum*, EC 3.2.1.11; Sigma-Aldrich, Darmstadt, Germany) at 37 °C for 16 h. The digested samples were centrifuged (5,000 × g, 5 min), and gDNA was isolated from the pellet using the NucleoSpin Food Mini kit (Macherey-Nagel, Düren, Germany) with minor adjustments. The pellet was suspended in 550 µL CF buffer with 10 µL RNase A solution (20 mg/mL; NEB, Ipswich, MA, USA), and DNA was eluted in 50 µL elution buffer.

For gDNA isolation from kefir liquid, 40 mL samples were taken and centrifuged (10,000 × g, 5 min). The cell pellet was resuspended in 2 mL sterile ¼ Ringer’s Solution (NaCl 2.25 g/L, KCl 0.105 g/L, NaHCO_3_ 0.05 g/L, CaCl_2_*6 H_2_O 0.12 g/L) and centrifuged again (10,000 × g, 2 min). The gDNA was isolated from the resulting cell pellet as described above.

DNA concentration and purity were measured with a NanoDrop© ND-1000 spectrophotometer (Thermo Fisher Scientific, Waltham, MA, USA). Sample integrity was checked on 1% (wt/vol) agarose gels containing ethidium bromide in 0.5X TAE buffer (20 mM Tris, 10 mM acetic acid, 0.5 mM EDTA). Samples were mixed with 6X DNA Loading Dye Purple (NEB, Ipswich, MA, USA) and compared to the 1 kb+ DNA ladder (NEB, Ipswich, MA, USA), with intact DNA detected above 10 kb.

Flow cytometry and fluorescence-activated cell sorting

### Sample preparation

To analyse microbial community dynamics in grains and liquid, standard fermentations with tap water and dried figs were performed in triplicate. Based on preliminary results, samples were collected at 0 h (substrate blank, before grain addition), 1 h, 2 h, 4 h, 6 h, 8 h, 24 h, 48 h, 72 h, 96 h, and 168 h. In addition to continuous OD_600_ measurements, the process parameters grain mass, pH, and TSS were recorded at the start and end of fermentation after 7 days to verify the proper course of fermentation. At each time point, 11 mL were withdrawn from the grain-free liquid phase; 1 mL was used for OD_600_ measurement, with the centrifuged supernatant as the blank, and the remaining 10 mL were centrifuged (10,000 × g, 5 min). Cell pellets were washed with 1 mL of ¼ Ringer’s solution, centrifuged again (5,000 × g, 2 min) and resuspended in 1 mL phosphate buffer (50 mM NaH_2_PO_4_/Na_2_HPO_4_, pH 6.5).

To analyse the microbial community in the grains, 100 mg of grain material were collected at each sampling time point, washed with 1 mL of ¼ Ringer’s solution, centrifuged (5,000 × g, 2 min), and resuspended in 1 mL of sodium phosphate buffer.

To degrade the EPS matrix, 10 µL dextranase solution (1:10 in sodium phosphate buffer) were added to each grain and liquid sample, gently mixed, and incubated at 28 °C for 18 h. Next, two cycles of 1 min of ultrasonication at 50 Hz (Bandelin Sonorex, Berlin, Germany), followed by 10 s of shaking with a vortex mixer, were applied to the samples to disaggregate remaining cell conglomerates.

For sample fixation, samples were transferred into glass tubes, and 1 mL PBS buffer (6 mM Na_2_HPO_4_, 1.8 mM NaH_2_PO_4_, 145 mM NaCl, pH 7.0) was added. Samples were centrifuged (3,200 × g, 10 min, 4 °C), and the supernatant was carefully removed. The pellets were resuspended in 1.5 mL PBS buffer and 0.5 mL of 8% (wt/vol) paraformaldehyde (PFA) solution, prepared as described by [66], and incubated for 30 min at room temperature. After a second centrifugation step (3,200 × g, 10 min, 4 °C), the supernatant was removed, and the cell pellets were resuspended in 2 mL 70% (vol/vol) ethanol. Samples were stored at -18 °C until further analysis.

### Cytometric analysis and cell sorting of selected subcommunities

For flow cytometric analysis, cellular DNA was stained with 0.24 µM 4′,6-diamidino-2-phenylindole (DAPI) according to the protocol by [66]. All samples were OD_700nm_ adjusted (0.5 cm path length) to 0.035. Measurements were carried out using a BD Influx v7 cell sorter (Becton, Dickinson and Company). DAPI-stained cells were excited with a 355 nm UV laser, and fluorescence emission was recorded through a 460 ± 25 nm band-pass filter. Forward scatter (FSC, 488 ± 5 nm band-pass filter), providing information related to cell size, and side scatter (SSC, 488 ± 5 nm band-pass filter, trigger), providing information to related to cell density, were excited with a 488 nm laser. Daily calibration of the cell sorter with beads and application of a biological standard (plate mCMC) was carried out as in [66]. 1 μm UV beads (BB Fluoresbrite, Polysciences, Warrington, PA, USA) were spiked additionally into each sample for comparability of all samples, respectively. For each sample, 200,000 cells were analysed and visualised in 2D dot plots displaying DAPI fluorescence intensity versus forward scatter (FSC) in the gate template excluding instrumental noise and beads. Gates were manually defined, and a unified gate template was subsequently applied across all samples to ensure comparability (see Figure 2A). Fluorescence-activated cell sorting (FACS) was performed for taxonomic profiling on selected subcommunities in a suitable replicate, based on their gate positions and relative abundance profiles. Per selected sub-community, 500,000 cells were sorted.

### Data Analysis

Flow cytometry data were analysed using the R packages flowCyBar and flowCHIC [56,67]. Before downstream analysis, cytometric data were imported into the latest FlowJo^TM^ V10 software package (Becton, Dickinson and Company, Franklin Lakes, NJ, USA), and a gate template was generated following the general approach described by [66]. Beads and instrumental noise were excluded, and cytometrically defined subcommunities were marked using ellipsoid gates. Non-overlapping gates were created for the most abundant subpopulations. Changes in subcommunity abundances in relation to temporal, spatial, abiotic, and biotic parameters were visualised using NMDS, boxplots, and heat maps generated with flowCyBar. Cytometric community structure was further compared using flowCHIC, which converts two-dimensional cytometric density plots to grayscale images and evaluates them based on the mean grayscale intensity of informative pixels. Similarities and differences in community composition were subsequently visualised by NMDS analysis.

### Amplicon sequencing and taxonomic profiling

Amplicon sequencing was used to characterise both the overall microbial communities of water kefir grains and fermentation liquid and selected flow-cytometrically sorted subcommunities. In both approaches, bacterial communities were analysed by amplification of the hypervariable V3–V4 region of the 16S rRNA gene using primers 341F (5′-CCT-ACG-GGN-GGC-WGC-AG-3′) and 805R (5′-GAC-TAC-HVG-GGT-ATC-TAA-TCC-3′), yielding an expected amplicon size of approximately 460 bp. Fungal communities were analysed by amplification of the ITS2 region between the 5.8S and 28S rRNA genes using primers ITS3 (5′-GCA-TCG-ATG-AAG-AAC-GCA-GC-3′) and ITS4 (5′-TCC-TCC-GCT-TAT-TGA-TAT-GC-3′), with an expected amplicon size of approximately 300–400 bp.

For profiling of the overall microbial communities, gDNA extracted from water kefir grains and fermentation liquid was submitted to Microsynth AG (Balgach, Switzerland) for Illumina amplicon sequencing and bioinformatic analysis. The standard bioinformatic workflow included read quality filtering, denoising, clustering and dereplication. Sequences were clustered into operational taxonomic units (OTUs), and taxonomic classification was performed using the RDP database [68] for bacterial 16S rRNA gene sequences and the UNITE database [69] for fungal ITS sequences. Representative sequences were additionally compared against the NCBI 16S/ITS databases using BLAST [70] and against the EzBioCloud database [71] to validate taxonomic assignments.

For taxonomic profiling of the sorted subcommunities, 70 µL of sterile 10% (wt/vol) Chelex 100 solution (Merck, Darmstadt, Germany) were added to each sample and mixed briefly. Samples were incubated for 45 min at 95 °C with shaking at 800 rpm in a Thermomixer 5436 (Eppendorf, Hamburg, Germany), followed by centrifugation at 7,000 × g for 5 min at 4 °C. Supernatants were transferred to sterile tubes while avoiding bead carryover, and DNA concentrations were quantified using a Qubit 3.0 fluorometer (Thermo Fisher Scientific, Waltham, MA, USA). The ZymoBIOMICS Microbial Community Standard (Zymo Research, Umkirch, Germany), comprising eight bacterial and two fungal strains, was processed in parallel as a process control.

Amplicons from the sorted subcommunities and the mock community were generated in-house using the primer pairs described above. For Illumina sequencing, primer sequences were combined with Illumina adapter sequences in a two-step PCR approach. The first PCR was performed in a total volume of 37.5 µL containing 18.75 µL 2x MyTaq Mix (BioCat, Heidelberg, Germany), 1.5 µL of each primer (0.4 µM), 1.5 µL BSA, and 1–10 µL DNA template, with PCR-grade water added to a total template volume of 14.25 µL. Amplification was performed using an S1000 thermocycler (Bio-Rad, Hercules, CA, USA) with an initial denaturation at 95 °C for 3 min, followed by 30 cycles of 95 °C for 30 s, 55 °C for 30 s, and 72 °C for 30 s, and a final extension at 72 °C for 5 min. PCR product concentrations were quantified using a Qubit 3.0 fluorometer, and amplicon sizes were verified by gel electrophoresis. Amplicons were subsequently submitted to Microsynth AG (Balgach, Switzerland) for further library processing and Illumina sequencing.

Raw sequence quality of the sorted subcommunities was assessed using FastQC v0.12.1 [72]. Primers and low-quality bases were removed using Cutadapt v5.1 [73], and amplicon sequence variants (ASVs) were inferred using the DADA2 pipeline v1.26.0 [74]. Taxonomic assignment was performed using SILVA v138 [75] for 16S rRNA gene sequences and the UNITE general FASTA release (2025-02-19) [69] for fungal ITS sequences. Representative sequences were additionally compared against the NCBI 16S/ITS databases using BLAST [70] and against EzBioCloud [71] to validate taxonomic assignments.

### Network-based comparison of water kefir communities

The microbial composition of the water kefir culture analysed in this study was compared with the 69 water kefir metagenomes reported by [32]. For this purpose, a consolidated presence–absence species profile was generated for our culture by integrating amplicon metagenomic sequencing, cultivation-based isolation, FACS-based sorting, and subsequent amplicon sequencing. This profile was combined with the species profiles of the published reference samples and used for network-based comparison. Species-level assignments were included when supported by at least one method with sufficient taxonomic resolution. Assignments that could not be resolved to a single species were retained as unresolved species-level labels.

Two complementary network representations were generated. First, a radial bipartite sample–species network was constructed to visualise species occurrence across the complete dataset. Individual water kefir samples were represented as one node set and microbial species as the second node set, with edges connecting each sample to the species detected in the respective culture. Sample nodes were coloured by country of origin, and species node size reflected prevalence across all samples. For unresolved taxonomic assignments, dashed edges were used to indicate links to the possible individual species identities.

Second, a sample–sample similarity network was constructed to quantify compositional overlap between water kefir communities. Pairwise taxonomic similarity between the present culture and each reference culture was calculated using the Jaccard index. A conservative Jaccard similarity (J_c_) was used as the primary metric, and only exact matches between unambiguous species-level labels were included. Unresolved assignments in the compiled profile were retained in the taxon set and therefore contributed to the union but were not counted as definitive matches to compatible single-species labels in the reference dataset. To assess the influence of unresolved assignments, a relaxed Jaccard similarity (J_r_) was additionally calculated as an upper-bound sensitivity estimate. In this approach, each unresolved assignment was allowed to match one compatible species in the reference sample if present. However, each unresolved assignment contributed at most one possible match and was not treated as a collapsed species group. Thus, if a reference sample contained multiple compatible species, only one was counted as a possible shared taxon, while the remaining species were retained as unmatched taxa in the union. The similarity network was generated using kefir samples as nodes and Jaccard similarities as edge weights.

Network construction, Jaccard similarity calculations, and network metrics were performed in Python v3.10 using NetworkX v3.4.2. The Jaccard similarity between samples was calculated as the number of shared taxa divided by the size of their union. Network figures were generated using custom visualisation tools based on the D3.js library.

### Grain growth studies

#### Technical setup

Kefir growth at the macroscopic level was monitored by time-lapse imaging. As the kefir structures appear white, their growth was quantified by measuring the number and increase of white pixels on a deep black background over time. For the technical implementation, a customised camera setup, including operating software, was developed and optimised. The aim was to capture images of growing water kefir under reproducible technical conditions (e.g., light and camera settings) with high temporal resolution. Information on the technical components, the internal setup diagram, and information on the control system can be found in the supplemental material.

#### Experimental design

Two experimental series were conducted. In the first series, single grains of different initial sizes (small, medium, large, and extra-large) were investigated to assess the general shape and reproducibility of pixel-based growth curves. Each condition was tested in biological triplicate in two independent 72-h experiments. This timeframe is based on a typical fermentation cycle in private households. For this series, only the total ROI was analysed, representing the absolute number of white pixels over time. The grain and liquid sub-ROIs were not evaluated, as grain motion and gas bubble formation in the larger (large, extra-large) samples led to unstable segmentation, while the signal in the smallest grains was largely dominated by growth in the liquid phase.

In the second series, samples containing two, four, or eight grains of the medium-size class were examined in biological triplicate. These experiments were repeated as three independent 72-hour experiments under identical conditions. In this series, all ROIs (total, grain, and liquid) were analysed, and derived growth parameters were calculated and statistically evaluated.

#### Cultivation medium

To monitor individual kefir grain growth with the camera setup, the fermentation batch size was reduced, and an optically clear water kefir medium (WKM) was prepared. The procedure was adapted from [76]. For the WKM, 18 g preservative-free dried figs (dmBio, Karlsruhe, Germany) were cut into pieces and extracted in 80 mL tap water in a baffled flask in a shaking incubator (180 rpm, 30 min, 37 °C). Larger pieces were removed by sieving and paper filtration, followed by centrifugation (5.000 × g, 20 min, 12 °C). The supernatant was sequentially filtered (0.45 µm, then 0.2 µm) to obtain a clear, sterile fruit extract, which was stored at -20 °C or used immediately. The final WKM contained 5% (wt/vol) refined sucrose (Südzucker, Mannheim, Germany), 1% (wt/vol) whole cane sugar (dmBio, Karlsruhe, Germany), and 10% (vol/vol) fig extract in tap water. The medium was sterile filtered (0.2 µm) and stored at 4 °C.

#### Pre-treatment of kefir grains and image acquisition

Kefir grains were pre-cultivated in a bulk fermentation for 24 h to ensure they were in an active stage. Grains were then sieved, and size-uniform grains were selected. To remove residual fermentation medium, grains were washed by agitation in sterile tap water on a platform shaker (40 mL, 40 rpm, 2 min). Washed grains were transferred to sterile crystallisation dishes, arranged individually or in groups, and supplemented with 15 mL WKM. The dishes were placed in the sample holder under the camera, and image acquisition was started, capturing two images (JPEG and DNG raw format) every 5 min over 72 h.

#### Image analysis and white pixel calculations from image-derived data

Image data (JPEG format) were first processed using the open-source software Fiji [77] before white pixel counts were evaluated. Images were converted into 8-bit black-and-white stacks. Image positions were corrected with the *Linear Stack Alignment with SIFT* module with default settings. Then, for each sample, three ROIs could be defined, depending on the experiment’s objectives: a total ROI, covering the entire visible sample area to capture the general kefir growth; a grain ROI, including only the grain structures and their growth behaviour; and a liquid ROI, representing the surrounding liquid phase without grains (Figure S2). The total ROI was applied as a constant circular mask encompassing the entire visible area of the crystallisation dish across all frames. The grain and liquid ROIs were defined separately based on the final image of each time series to capture the final grain dimensions at the end of the experiment. To ensure that grain movement or morphological changes during growth did not introduce segmentation errors, the entire image stack was visually inspected and manually verified for consistency of the ROIs across time. Because the grains and liquid ROIs were defined independently rather than derived by subtraction, the total ROI does not represent the exact sum of the two sub-ROIs but serves as an independent measure of the overall growth signal. ROI sets were created and saved separately for each experiment. For all experiments, only the total ROI size was identical, corresponding to the maximum achievable value of white pixels (430,542 px). The ROI set was applied to the corresponding image stack, and the mean grey values (*mean_grey_*) and ROI areas (*area_ROI_*) were determined. Then, the relative percentage of white pixels (*px_w,rel_)* and the absolute number of all white pixels (*px_w_*) were calculated (for equations, see supplemental material). Finally, the absolute numbers of pixels for overall growth, grain growth, and liquid-phase growth were displayed over time and used for further analyses.

Data analysis was performed using custom Python scripts executed in the Thonny Python IDE v4.1.1 with Python v3.10.11, employing the libraries *numpy*, *pandas*, and *scipy.stats*. Statistical analysis, data visualisation, and plotting were performed using OriginPro 2024 v10.1.0.178. Biological replicates were generated within each experiment, and experiments were independently repeated on three different starting days. For statistical analysis, measurements from biological replicates were averaged within each experiment to avoid pseudo-replication, yielding *n = 3* independent experiments per tested condition.

### Structural analysis of water kefir grains

#### Stereomicroscopy of intact grains

For documenting the macroscopic morphology, kefir grains of different sizes were selected, rinsed, and the adhering water was carefully removed with a lint-free cloth. Pictures of the grains were taken with the Stereomicroscope MZ10 F (Leica Microsystems, Wetzlar, Germany) and the ProgRes SpeedXT core5 camera (Jenoptik, Jena, Germany).

#### Scanning electron microscopy of intact grains

Two medium-sized grains were sampled from the batch and rinsed with tap water. One grain was left intact, while the other one was cut in half with a sterile scalpel. The samples were dehydrated in a graded series of ethanol-water mixtures (70%, 80%, 90%, 96%, 100%) and critical-point dried with liquid carbon dioxide using a CPD 030 critical-point dryer (BAL-TEC, Balzers, Liechtenstein). Then, the samples were attached to sample carriers with conductive double-sided adhesive carbon tab and coated with gold in an Emitech K550 sputter coater (Quorum Technologies, Laughton, United Kingdom). The samples were examined using a Zeiss Supra 40VP scanning electron microscope (Carl Zeiss Microscopy, Jena, Germany) at an accelerating voltage of 5 kV.

#### Fixation and embedding

Medium-sized grains were sampled from the batch, rinsed with tap water and fixed in modified Karnovsky’s fixative (2% (wt/vol) paraformaldehyde, 2% (wt/vol) glutaraldehyde in 4-(2-hydroxyethyl)-1-piperazine-ethanesulfonic acid (HEPES) buffer (50 mM, pH 7.4)) for 1 h at room temperature and 24 h at 4 °C. Fixed grains were cut into quarters, washed once in HEPES buffer (100 mM, pH 7.4) and three times in water (5 min each). Then, the grains were stained on ice for 30 min with a 1:1 mixture of 3% (wt/vol) K_4_[Fe(CN)_6_] in 4 mM CaCl_2_*2H_2_O and 4% (wt/vol) OsO_4_, and washed three times with water (5 min each). Samples were incubated in 1% (wt/vol) thiocarbohydrazide for 20 min, washed five times in water (5 min each), and stained with 2% (wt/vol) OsO_4_ for 30 min on ice. After three further washes (5 min each), samples were stained with 1% (wt/vol) uranyl acetate overnight at 4 °C and washed three times the next day. Dehydration was carried out in a graded series of ethanol in water (30%, 50%, 70%, 90%, 96%, 100%), followed by infiltration with Araldite 502/ Epon-812 epoxy resin (Science Services, Munich, Germany) in a graded series in ethanol (25%, 50%, 75%, 100%). The final polymerisation was performed for 48 h at 60 °C.

#### Cross-sectioning, scanning electron microscopy, and transmission electron microscopy

Sections of 1 µm thickness were cut from embedded grain samples using an EM UC6 ultra-microtome (Leica Microsystems, Wetzlar, Germany), with 50–150 µm spacing to access different depths of the grains. The sections were placed on silicon wafers and sputter-coated with 10 nm carbon using a CCU-010 HV sputter coater (Safematic, Zizers, Switzerland). Samples were examined with a ZEISS Crossbeam 550 scanning electron microscope (Carl Zeiss Microscopy, Jena, Germany) at an accelerating voltage of 8 kV, using a backscattered electron (BSE) detector.

Two samples previously analysed by SEM were selected for subsequent TEM analysis. Ultrathin sections of 70–100 nm thickness were prepared and placed on TEM grids (300 mesh, hexagonal, copper, with a Formvar resin and carbon support film; Science Services, Munich, Germany). The sections were stained with 4% (wt/vol) uranyl acetate for 10 min, washed three times with water (1 min each), incubated in 0.4% (wt/vol) lead citrate for 90 s, washed three times again, and dried with filter paper. Imaging was performed with a Jeol JEM-1400Plus transmission electron microscope (JEOL, Akishima, Japan) at 80 kV accelerating voltage.

## Data Availability

All cytometric raw data, including the .fcs files and the FlowJo workspace with the gate template, are stored in the Zenodo database under: https://zenodo.org/records/21357475 (DOI: 10.5281/zenodo.21357475).

The raw 16S rRNA and ITS2 amplicon sequencing data generated in this study have been deposited in the NCBI Sequence Read Archive (SRA) under accession number PRJNAxxx [*The raw amplicon sequencing data will be deposited in the NCBI Sequence Read Archive (SRA), and the corresponding accession number will be provided during the review process.]*.

The time-lapse videos generated in this study will be made available to reviewers via a private link during the peer-review process and will be made publicly accessible upon publication of the peer-reviewed manuscript.

## ACKNOWLEDGEMENTS

This work was supported by the German Research Foundation (DFG) within the priority program SPP 2389 “Emergent Functions of Bacterial Multicellularity” (grants 504017689 (to T.M.) and 503905203 (to S.M.)) and by a Maria Reiche Doctoral Fellowship of the TUD Dresden University of Technology Graduate Academy (to P.B.). The authors would like to thank Susanne Kretschmar for TEM sample preparation, Alice Ludewig for coordinating TEM work and performing SEM imaging of cross-sections, Thomas Kurth for TEM imaging of cross-sections, Luca Bertinetti for consultation on BSE imaging and Markus Günther for SEM imaging of kefir grains. The authors would also like to acknowledge the support of Daniel Rentzsch in constructing the camera hardware and programming the software, and Isin Kosemen for her support with data visualisation.

## REFERENCES

1. Szathmáry E, Smith JM. The major evolutionary transitions. Nature. 1995;374: 227–232. doi:10.1038/374227a0

2. Grosberg RK, Strathmann RR. The Evolution of Multicellularity: A Minor Major Transition? Annu Rev Ecol Evol Syst. 2007;38: 621–654. doi:10.1146/annurev.ecolsys.36.102403.114735

3. Knoll AH. The Multiple Origins of Complex Multicellularity. Annu Rev Earth Planet Sci. 2011;39: 217–239. doi:10.1146/annurev.earth.031208.100209

4. Parfrey LW, Lahr DJG. Multicellularity arose several times in the evolution of eukaryotes (Response to DOI 10.1002/bies.201100187). BioEssays. 2013;35: 339–347. doi:10.1002/bies.201200143

5. Bonner JT. The origins of multicellularity. Integr Biol. 1998;1: 27–36. doi:10.1002/(SICI)1520-6602(1998)1:1<27::AID-INBI4>3.0.CO;2-6

6. Michod RE. Evolution of individuality during the transition from unicellular to multicellular life. Proc Natl Acad Sci USA. 2007;104: 8613–8618. doi:10.1073/pnas.0701489104

7. Tarnita CE, Taubes CH, Nowak MA. Evolutionary construction by staying together and coming together. Journal of Theoretical Biology. 2013;320: 10–22. doi:10.1016/j.jtbi.2012.11.022

8. Herron MD, Borin JM, Boswell JC, Walker J, Chen I-CK, Knox CA, et al. De novo origins of multicellularity in response to predation. Sci Rep. 2019;9: 2328. doi:10.1038/s41598-019-39558-8

9. Strassmann JE, Zhu Y, Queller DC. Altruism and social cheating in the social amoeba Dictyostelium discoideum. Nature. 2000;408: 965–967. doi:10.1038/35050087

10. West SA, Fisher RM, Gardner A, Kiers ET. Major evolutionary transitions in individuality. Proc Natl Acad Sci U S A. 2015;112: 10112–10119. doi:10.1073/pnas.1421402112

11. Lyons NA, Kolter R. On the evolution of bacterial multicellularity. Current Opinion in Microbiology. 2015;24: 21–28. doi:10.1016/j.mib.2014.12.007

12. Claessen D, Rozen DE, Kuipers OP, Søgaard-Andersen L, Van Wezel GP. Bacterial solutions to multicellularity: a tale of biofilms, filaments and fruiting bodies. Nat Rev Microbiol. 2014;12: 115–124. doi:10.1038/nrmicro3178

13. Flores E, Herrero A. Compartmentalized function through cell differentiation in filamentous cyanobacteria. Nat Rev Microbiol. 2010;8: 39–50. doi:10.1038/nrmicro2242

14. Nadell CD, Drescher K, Foster KR. Spatial structure, cooperation and competition in biofilms. Nat Rev Microbiol. 2016;14: 589–600. doi:10.1038/nrmicro.2016.84

15. Kolter R, Greenberg EP. The superficial life of microbes. Nature. 2006;441: 300–302. doi:10.1038/441300a

16. Flemming H-C, Wingender J. The biofilm matrix. Nat Rev Microbiol. 2010;8: 623– 633. doi:10.1038/nrmicro2415

17. O’Toole G, Kaplan HB, Kolter R. Biofilm Formation as Microbial Development. Annu Rev Microbiol. 2000;54: 49–79. doi:10.1146/annurev.micro.54.1.49

18. Stewart PS, Franklin MJ. Physiological heterogeneity in biofilms. Nat Rev Microbiol. 2008;6: 199–210. doi:10.1038/nrmicro1838

19. Elias S, Banin E. Multi-species biofilms: living with friendly neighbors. FEMS Microbiol Rev. 2012;36: 990–1004. doi:10.1111/j.1574-6976.2012.00325.x

20. Kolenbrander PE, Palmer RJ, Periasamy S, Jakubovics NS. Oral multispecies biofilm development and the key role of cell–cell distance. Nat Rev Microbiol. 2010;8: 471–480. doi:10.1038/nrmicro2381

21. Bowen WH, Burne RA, Wu H, Koo H. Oral Biofilms: Pathogens, Matrix, and Polymicrobial Interactions in Microenvironments. Trends in Microbiology. 2018;26: 229–242. doi:10.1016/j.tim.2017.09.008

22. Mark Welch JL, Ramírez-Puebla ST, Borisy GG. Oral Microbiome Geography: Micron-Scale Habitat and Niche. Cell Host & Microbe. 2020;28: 160–168. doi:10.1016/j.chom.2020.07.009

23. Harriott MM, Noverr MC. Importance of Candida–bacterial polymicrobial biofilms in disease. Trends in Microbiology. 2011;19: 557–563. doi:10.1016/j.tim.2011.07.004

24. Korgaonkar A, Trivedi U, Rumbaugh KP, Whiteley M. Community surveillance enhances *Pseudomonas aeruginosa* virulence during polymicrobial infection. Proc Natl Acad Sci USA. 2013;110: 1059–1064. doi:10.1073/pnas.1214550110

25. Donaldson GP, Lee SM, Mazmanian SK. Gut biogeography of the bacterial microbiota. Nat Rev Microbiol. 2016;14: 20–32. doi:10.1038/nrmicro3552

26. Stams AJM, Plugge CM. Electron transfer in syntrophic communities of anaerobic bacteria and archaea. Nat Rev Microbiol. 2009;7: 568–577. doi:10.1038/nrmicro2166

27. Fiorda FA, Melo Pereira GV, Thomaz-Soccol V, Rakshit SK, Pagnoncelli MGB, Vandenberghe LP de S, et al. Microbiological, biochemical, and functional aspects of sugary kefir fermentation - A review. Food Microbiology. 2017;66: 86–95. doi:10.1016/j.fm.2017.04.004

28. Laureys D, De Vuyst L. Microbial species diversity, community dynamics, and metabolite kinetics of water Kefir fermentation. Applied and Environmental Microbiology. 2014;80: 2564–2572. doi:10.1128/AEM.03978-13

29. Gulitz A, Stadie J, Wenning M, Ehrmann MA, Vogel RF. The microbial diversity of water kefir. International Journal of Food Microbiology. 2011;151: 284–288. doi:10.1016/j.ijfoodmicro.2011.09.016

30. Patel SH, Tan JP, Börner RA, Zhang SJ, Priour S, Lima A, et al. A temporal view of the water kefir microbiota and flavour attributes. Innovative Food Science and Emerging Technologies. 2022;80. doi:10.1016/j.ifset.2022.103084

31. Zannini E, Lynch KM, Nyhan L, Sahin AW, O’ Riordan P, Luk D, et al. Influence of Substrate on the Fermentation Characteristics and Culture-Dependent Microbial Composition of Water Kefir. Fermentation. 2022;9: 28. doi:10.3390/fermentation9010028

32. Breselge S, Skibinska I, Yin X, Brennan L, Kilcawley K, Cotter PD. The core microbiomes and associated metabolic potential of water kefir as revealed by pan multi-omics. Communications biology. 2025;8: 415. doi:10.1038/s42003-025-07808-3

33. Laureys D, Leroy F, Vandamme P, Vuyst L. Backslopping Time, Rinsing of the Grains During Backslopping, and Incubation Temperature Influence the Water Kefir Fermentation Process. Frontiers in Microbiology. 2022;13: 871550. doi:10.3389/fmicb.2022.871550

34. Pidoux M, Brillouet JM, Quemener B. Characterization of the polysaccharides from a Lactobacillus brevis and from sugary kefir grains. Biotechnology Letters. 1988;10: 415–420.

35. Lynch KM, Wilkinson S, Daenen L, Arendt EK. An update on water kefir: Microbiology, composition and production. International Journal of Food Microbiology. 2021;345: 109–128. doi:10.1016/j.ijfoodmicro.2021.109128

36. Wang X, Li W, Xu M, Tian J, Li W. The Microbial Diversity and Biofilm-Forming Characteristic of Two Traditional Tibetan Kefir Grains. Foods (Basel, Switzerland). 2021;11. doi:10.3390/foods11010012

37. Laureys D, Van Jean A, Dumont J, De Vuyst L. Investigation of the instability and low water kefir grain growth during an industrial water kefir fermentation process. Applied Microbiology and Biotechnology. 2017;101: 2811–2819. doi:10.1007/s00253-016-8084-5

38. Fels L, Jakob F, Vogel RF, Wefers D. Structural characterization of the exopolysaccharides from water kefir. Carbohydrate Polymers. 2018;189: 296–303. doi:10.1016/j.carbpol.2018.02.037

39. Davidovic S, Miljkovic M, Antonovic D, Rajilic-Stojanovic M, Dimitrijevic-Brankovic S. Water Kefir grain as a source of potent dextran producing lactic acid bacteria. Hem Ind. 2015;69: 595–604. doi:10.2298/HEMIND140925083D

40. Bechtner J, Hassler V, Wefers D, Vogel RF, Jakob F. Insights into extracellular dextran formation by Liquorilactobacillus nagelii TMW 1.1827 using secretomes obtained in the presence or absence of sucrose. Enzyme and Microbial Technology. 2021;143. doi:10.1016/j.enzmictec.2020.109724

41. Tan LL, Ngiam JJ, Sim ESZ, Conway PL, Loo SCJ. Liquorilactobacillus satsumensis from water kefir yields α-glucan polysaccharides with prebiotic and synbiotic qualities. Carbohydrate Polymers. 2022;290. doi:10.1016/j.carbpol.2022.119515

42. Pendón MD, Bengoa AA, Iraporda C, Medrano M, Garrote GL, Abraham AG. Water kefir: Factors affecting grain growth and health-promoting properties of the fermented beverage. Journal of Applied Microbiology. 2022;133: 162–180. doi:10.1111/jam.15385

43. Magalhães KT, M Pereira GV de, Dias DR, Schwan RF. Microbial communities and chemical changes during fermentation of sugary Brazilian kefir. World journal of microbiology & biotechnology. 2010;26: 1241–1250. doi:10.1007/s11274-009-0294-x

44. Marsh AJ, O’Sullivan O, Hill C, Ross RP, Cotter PD. Sequence-based analysis of the microbial composition of water kefir from multiple sources. FEMS Microbiology Letters. 2013;348: 79–85. doi:10.1111/1574-6968.12248

45. Martínez-Torres A, Gutiérrez-Ambrocio S, Heredia-del-Orbe P, Villa-Tanaca L, Hernández-Rodríguez C. Inferring the role of microorganisms in water kefir fermentations. International Journal of Food Science and Technology. 2017;52: 559–571. doi:10.1111/ijfs.13312

46. Bottazzi V, Bianchi F. A Note on Scanning Electron Microscopy of Micro-organisms associated with the Kefir Granule. Journal of Applied Bacteriology. 1980;48.

47. Neve H, Heller KJ. The microflora of water kefir: A glance by scanning electron microscopy. Kieler Milchwirtschaftliche Forschungsberichte. 2002;54: 337–349.

48. Laureys D, Leroy F, Hauffman T, Raes M, Aerts M, Vandamme P, et al. The Type and Concentration of Inoculum and Substrate as Well as the Presence of Oxygen Impact the Water Kefir Fermentation Process. Frontiers in Microbiology. 2021;12. doi:10.3389/fmicb.2021.628599

49. Moinas M, Horisberger M, Bauer H. The structural organization of the Tibi grain as revealed by light, scanning and transmission microscopy. Archives of Microbiology. 1980; 157–161. doi:10.1007/BF00406153

50. Laureys D, Aerts M, Vandamme P, De Vuyst L. Oxygen and diverse nutrients influence the water kefir fermentation process. Food Microbiology. 2018;73: 351–361. doi:10.1016/j.fm.2018.02.007

51. Laureys D, Aerts M, Vandamme P, Vuyst L. The Buffer Capacity and Calcium Concentration of Water Influence the Microbial Species Diversity, Grain Growth, and Metabolite Production During Water Kefir Fermentation. Frontiers in Microbiology. 2019;10: 2876. doi:10.3389/fmicb.2019.02876

52. Wang SY, Chen KN, Lo YM, Chiang ML, Chen HC, Liu JR, et al. Investigation of microorganisms involved in biosynthesis of the kefir grain. Food Microbiology. 2012;32: 274–285. doi:10.1016/j.fm.2012.07.001

53. Dong J, Liu B, Jiang T, Liu Y, Chen L. The biofilm hypothesis: The formation mechanism of Tibetan kefir grains. International Journal of Dairy Technology. 2018;71: 44–50. doi:10.1111/1471-0307.12473

54. Han X, Zhang LJ, Wu HY, Wu YF, Zhao SN. Investigation of microorganisms involved in kefir biofilm formation. Antonie van Leeuwenhoek, International Journal of General and Molecular Microbiology. 2018;111: 2361–2370. doi:10.1007/s10482-018-1125-6

55. Xu D, Fels L, Wefers D, Behr J, Jakob F, Vogel RF. Lactobacillus hordei dextrans induce Saccharomyces cerevisiae aggregation and network formation on hydrophilic surfaces. International Journal of Biological Macromolecules. 2018;115: 236–242. doi:10.1016/j.ijbiomac.2018.04.068

56. Koch C, Fetzer I, Schmidt T, Harms H, Müller S. Monitoring functions in managed microbial systems by cytometric bar coding. Environmental science & technology. 2013;47: 1753–1760. doi:10.1021/es3041048

57. Breselge S, De Paula Dias Moreira L, Skibinska I, Yin X, Brennan L, Kilcawley K, et al. Water kefir multi-omics reveals functional redundancies despite taxonomic differences and the underappreciated contribution of yeast. npj Sci Food. 2025;9: 265. doi:10.1038/s41538-025-00624-3

58. Michielsen S, Vercelli GT, Cordero OX, Bachmann H. Spatially structured microbial consortia and their role in food fermentations. Current Opinion in Biotechnology. 2024;87: 103102. doi:10.1016/j.copbio.2024.103102

59. Pidoux M. The microbial flora of sugary kefir grain (the gingerbeer plant): biosynthesis of the grain from Lactobacillus hilgardii producing a polysaccharide gel. MIRCEN Journal of Apllied Microbiology and Biotechnology. 1989;5: 223–238.

60. Marshall VM, Cole WM, Brooker BE. Observations on the structure of kefir grains and the distribution of the microflora. Journal of Applied Bacteriology. 1984; 491–497.

61. Sutherland IW. The biofilm matrix - an immobilized but dynamic microbial environment. TRENDS in Microbiology. 2001; 222–227.

62. Simões M, Simões LC, Vieira MJ. A review of current and emergent biofilm control strategies. LWT - Food Science and Technology. 2010; 573–583. doi:10.1016/j.lwt.2009.12.008

63. Henderson A, Del Panta A, Schubert OT, Mitri S, Van Vliet S. Disentangling the feedback loops driving spatial patterning in microbial communities. npj Biofilms Microbiomes. 2025;11: 32. doi:10.1038/s41522-025-00666-1

64. Cai Y-M. Non-surface Attached Bacterial Aggregates: A Ubiquitous Third Lifestyle. Front Microbiol. 2020;11: 557035. doi:10.3389/fmicb.2020.557035

65. Espinoza Miranda SS, Abbaszade G, Hess WR, Drescher K, Saliba A-E, Zaburdaev V, et al. Resolving spatiotemporal dynamics in bacterial multicellular populations: approaches and challenges. Van Der Meer JR, editor. Microbiol Mol Biol Rev. 2025;89: e00138–24. doi:10.1128/mmbr.00138-24

66. Cichocki N, Hübschmann T, Schattenberg F, Kerckhof F-M, Overmann J, Müller S. Bacterial mock communities as standards for reproducible cytometric microbiome analysis. Nature Protocols. 2020;15: 2788–2812. doi:10.1038/s41596-020-0362-0

67. Koch C, Fetzer I, Harms H, Müller S. CHIC-an automated approach for the detection of dynamic variations in complex microbial communities. Cytometry Part A : the journal of the International Society for Analytical Cytology. 2013;83: 561–567. doi:10.1002/cyto.a.22286

68. Maidak BL, Olsen GJ, Larsen N, Overbeek R, McCaughey MJ, Woese CR. The RDP (Ribosomal Database Project). Nucleic Acids Res. 1997;25: 109–111. doi:10.1093/nar/25.1.109

69. Abarenkov K, Zirk A, Piirmann T, Pöhönen R, Ivanov F, Nilsson RH, et al. UNITE general FASTA release for Fungi. UNITE Community; 2025. doi:10.15156/BIO/3301229

70. Altschul SF, Gish W, Miller W, Myers EW, Lipman DJ. Basic local alignment search tool. Journal of Molecular Biology. 1990;215: 403–410. doi:DOI: 10.1016/S0022-2836(05)80360-2

71. Chalita M, Kim YO, Park S, Oh H-S, Cho JH, Moon J. EzBioCloud: a genome-driven database and platform for microbiome identification and discovery. InInternational journal of systematic and evolutionary microbiology. 2024;74. doi:10.1099/ijsem.0.006421

72. Andrews S. FastQC: A Quality Control Tool for High Throughput Sequence Data. In: Babraham Bioinformatics [Internet]. Available: https://www.bioinformatics.babraham.ac.uk/projects/fastqc/

73. Martin M. Cutadapt removes adapter sequences from high-throughput sequencing reads. EMBnet j. 2011;17: 10. doi:10.14806/ej.17.1.200

74. Callahan BJ, McMurdie PJ, Rosen MJ, Han AW, Johnson AJA, Holmes SP. DADA2: High-resolution sample inference from Illumina amplicon data. Nat Methods. 2016;13: 581–583. doi:10.1038/nmeth.3869

75. Quast C, Pruesse E, Yilmaz P, Gerken J, Schweer T, Yarza P, et al. The SILVA ribosomal RNA gene database project: improved data processing and web-based tools. Nucleic Acids Research. 2012;41: 590–596. doi:10.1093/nar/gks1219

76. Stadie J, Gulitz A, Ehrmann MA, Vogel RF. Metabolic activity and symbiotic interactions of lactic acid bacteria and yeasts isolated from water kefir. Food Microbiology. 2013;35: 92–98. doi:10.1016/j.fm.2013.03.009

77. Schindelin J, Arganda-Carreras I, Frise E, Kaynig V, Longair M, Pietzsch T. Fiji: an open-source platform for biological-image analysis. Nat Methods. 2012;9: 676–682. doi:10.1038/nmeth.2019

