## Supplemental Material for "Water Kefir as a Model for Aggregative Development of Microbial Multicellularity within a Multispecies Biofilm"

### Supplemental methods

#### Camera setup for kefir growth studies

The hardware components of the camera setup are listed in Table S1, and a schematic diagram is shown in Figure S1. All components were permanently mounted in stable, light- and dust-proof housing. A Raspberry Pi High-Quality Camera with a telephoto lens was connected to a Raspberry Pi 4B computer via the Camera Serial Interface and mounted 72 cm above the samples at a 90° angle to maintain a constant focal plane across the recording area.

Illumination was provided by light-emitting diode (LED) strips mounted on aluminium profiles. To achieve uniform and variable illumination, one LED panel consisting of 18 LEDs, arranged in three rows 4 cm apart, was mounted on two opposite sides of the samples using ball joint arms, outside the camera's field of view. The lamps were positioned 55 cm apart at a 45° angle to the sample, ensuring that reflections only occurred at the edges of the sample vessels. The LED strips were operated at full power using a controller. The circuit was created using a transistor, a 10 Ω resistor, and the general-purpose input/output (GPIO) of the connected Raspberry Pi. When selecting the controller, attention was paid to having a sufficiently high repetition frequency, and each LED strip was fitted with a diffusion attachment.

Reflections were minimised using polarisation filters. One side of each LED panel was covered with linear polarisation film, and a polarisation filter was mounted in front of the lens. The filter was rotated and fixed to achieve maximum extinction. The procedure was repeated for the second LED panel, aligning the polarising film to further reduce reflections in the sample vessels.

Crystallisation dishes were used as sample vessels and placed in a custom-made holder positioned 8 cm from the background. The holder minimised light beneath the samples, creating a deep black background, and all components were painted matte black where possible to reduce reflections. It accommodated nine dishes, allowing three fermentation conditions to be tested in triplicate. The holder was equipped with four black-and-white markers for subsequent image stabilisation and for measuring size.

The camera and LED panels were controlled via a Python script using the Picamera2 library. This program was used to configure the camera's exposure and gain parameters, as well as the image capture frequency and number of captured images. The focus and aperture were set directly on the lens. An f/6 aperture was selected. The exposure time was set to $2*10^6$ µs and the gain to a factor of 3.

The program sequence was designed to switch on the lamps shortly before image acquisition. First, a JPEG-format image was captured, followed by a DNG (RAW)-format image. After saving both images, the lamps were switched off until the next acquisition cycle. This was done to prevent the lamps from emitting heat into the environment through continuous illumination, thereby altering growth conditions. To monitor the growth conditions, the setup could be additionally equipped with an optional sensor for environmental factors such as temperature and humidity.

Table S1. Hardware components and specifications for the camera station used in water kefir growth studies

| Component | Specification | Manufacturer |
|---|---|---|
| Aluminium profiles | L-profile, flat | Grosfillex, Arbent |
| Diffusion attachment | / | Paulmann, Springe-Voelksen |
| Wooden shelf | Model "IVAR", dimensions 89x50x148 cm | IKEA, Munich, Germany |
| Crystallisation dishes | Volume: 20 ml, diameter: 40 mm, material: borosilicate glass | Merck, Darmstadt, Germany |
| LED strips | 24 V, cool white 600 K, 7500 lm | Shenzhen JoyLite, Guangdong, China |
| LED controller | 750 Hz flicker-free, 12–36 V | iluminize GmbH, Issum-Sevelen, Germany |
| Magic arms | 10 inches | SmallRig, Mongkok, Hongkong |
| Polarisation film | Typ ST-38-20, linear 90° | Screen-Tech, Hohenaspe, Germany |
| Raspberry Pi ® 4 Model B | 8 GB RAM, operating system: Raspberry Pi OS | Raspberry Pi Foundation, Wales, United Kingdom |
| Raspberry Pi ® High Quality Camera | Sensor: Sony IMX477 (12.3 Megapixels), C/CS-Mount | Raspberry Pi Foundation, Wales, United Kingdom |
| Telephoto lens | Focal length: 25 mm, C-Mount, aperture: f/1,4-16 | Berrybase, Berlin, Germany |
| Transistor | BC547 | Conrad, Hirschau, Germany |
| Resistor | 10 Ω | Conrad, Hirschau, Germany |
| Circular polarisation filter | Kaesemann High Transmission MRC Nano Master 82 mm | Schneider-Kreuznach, Bad Kreuznach, Germany |
| Foil curtain | PVC, black, material strength 1 mm, grammage 1270 $g/m^2$ | Heissner GmbH, Lauterbach, Germany |
| Spray paint | Matte, deep black, RAL 9005 | Maston One, Veikkola, Finland |

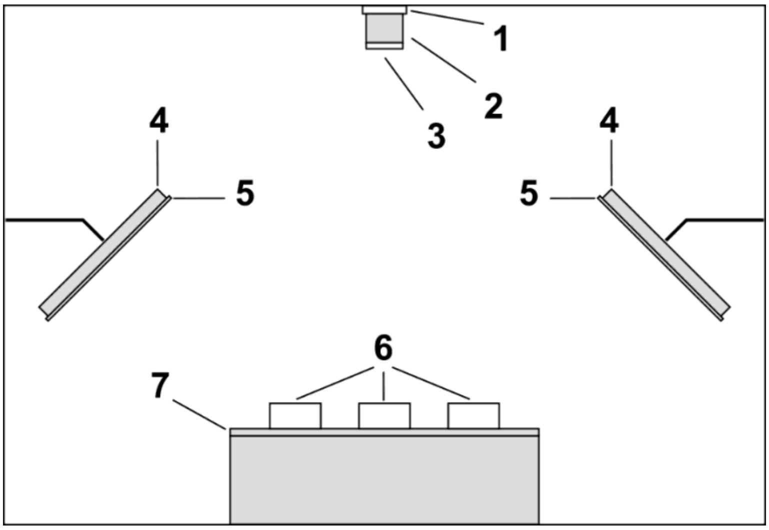


Figure S1. Schematic illustration of the internal camera setup with (**1**) Raspberry Pi ® high-quality camera, (**2**) telephoto lens, (**3**) circular polarisation filter, (**4**) LED panels, (**5**) polarisation film, (**6**) crystallisation dishes, and (**7**) sample holder. The illustration is not to scale.

### Determination of the region of interest during image analysis

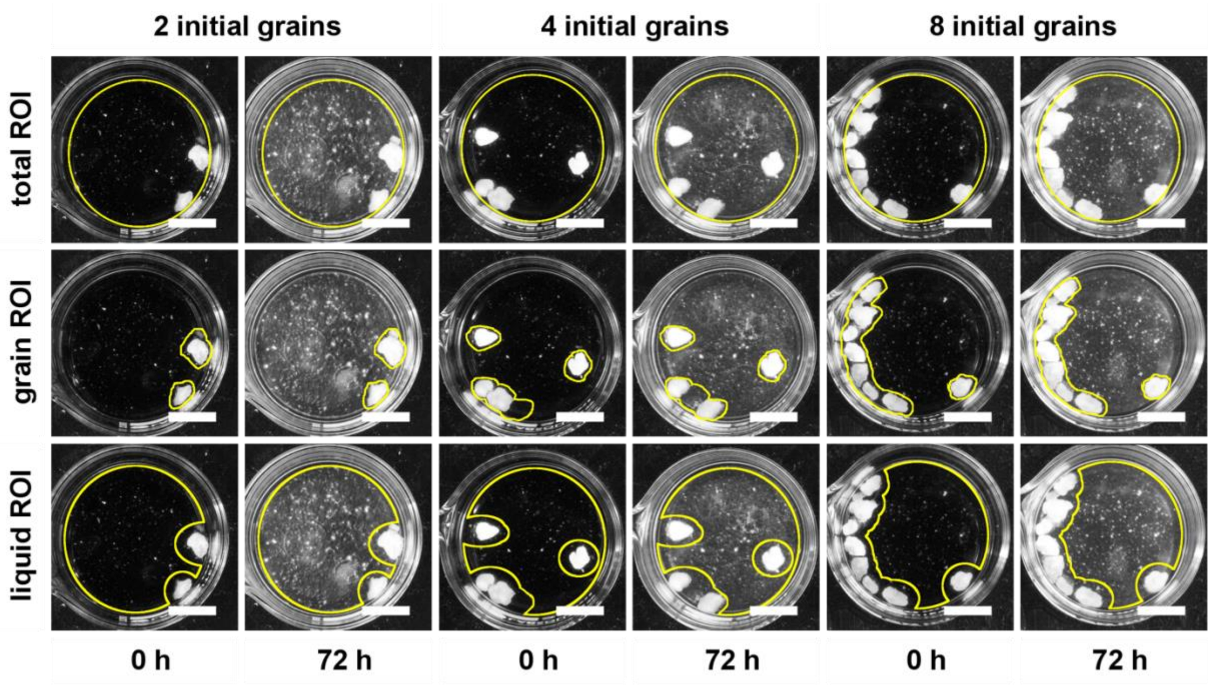


Figure S2. Exemplary form and placement of regions of interest (ROI) used for image analysis. Representative images showing the total ROI, grain ROI, and liquid ROI for samples containing 2, 4, and 8 initial grains at the beginning of the experiment and after 72 h. Scale bars represent 10 mm.

### Calculation of white pixels from image-derived data

To calculate white pixel values from image data, three biological replicates per treatment group and experimental day were averaged. The number of white pixels was estimated by assuming that the brightness in each area reflected the ratio of black to white pixels. Based on the possible brightness values of 8-bit grayscale images, ranging from 0 for pure black to 255 for pure white, and the measured $mean_{grey}$ value, the relative percentage of white pixels $px_{w,rel}$ was calculated. Based on the region of interest (ROI) area given in pixels, the absolute number of all white pixels $px_w$ could be calculated.

$$px_{w,rel}[\%] = mean_{grey} * \frac{100}{255} \tag{1.1}$$

$$px_w[absolute] = \frac{px_{w,rel}}{100\%} * area_{ROI} \tag{1.2}$$

### Calculation of growth parameter from image-derived data

Due to movement artifacts and bubble formation, especially in experiments with large and extra-large grains, sub-ROI analysis was not feasible in the size series. Therefore, sub-ROI analyses were performed only in the grain number series using 2, 4, and 8 medium-sized starter grains. This setup provided the most stable image series and the most reproducible signal dynamics, enabling reliable subsequent calculations of growth parameters.

The minimum and maximum white pixel counts were computed as the means of all values recorded during the first and last 60 min, respectively, to account for any potential fluctuations caused by technical variation. The total increase in white pixels $\Delta px_w$ was then determined as the difference between these two mean values, representing the absolute increase in white pixels over the course of the 72-h observation period.

$$\Delta px_w = \overline{px_{w,min}} - \overline{px_{w,max}} \tag{2.1}$$

$$\Delta px_w = \frac{1}{n}\sum_{t=t_{end}-60min}^{t_{end}} px_w(t) - \frac{1}{n}\sum_{t=t_0}^{t_0+60min} px_w(t) \tag{2.2}$$

To identify key transitions within the growth process, two thresholds were defined. The estimated transition from the lag phase to the exponential phase was set at 5% of total growth (lower threshold, $px_{w,5\%}$), and from the exponential to the stationary phase at 70% of total growth (upper threshold, $px_{w,70\%}$). The 5% and 70% thresholds were determined from multiple pre-experiments to provide consistent reference points that approximately correspond to the beginning and end of the exponential growth phase in the total ROI. Accordingly, these thresholds do not represent biologically defined transition points but rather standardised, empirically selected reference points for quantitative comparison between experiments.

$$lower\ threshold\ px_{w,5\%} = \Delta px_w * 0.05 + \overline{px_{w,min}} \tag{3.1}$$

$$upper\ threshold\ px_{w,70\%} = \Delta px_w * 0.70 + \overline{px_{w,min}} \tag{3.2}$$

To assign corresponding time stamps to the lower and upper thresholds, the five smallest white pixel values that exceeded the lower threshold were selected. The earliest of these was then chosen to mark the onset of exponential growth. Similarly, the five largest values below the upper threshold were extracted, and the latest timestamp was selected to indicate the end of exponential growth and the start of the stationary phase.

In analogy to the conventional definition of generation time as the period required for a microbial population to double during the exponential phase, we here define the pixel-doubling time *D* as the period required for the number of white pixels to double. First, the natural logarithm of each time series $px_w(t)$ was calculated, and then the curve was slightly smoothed using the Savitzky–Golay method with a second-order filter and a window width of 20 points. The temporal derivative of this curve was then determined. The maximum value $r_{max}$ was then identified within the exponential signal range, while excluding the edge points and early noise. The pixel-doubling time *D* could then be calculated from this.

Exponential growth model:

$$px_w(t) = px_{w,0}e^{rt} \tag{4.1}$$

$$\ln px_w(t) = \ln px_{w,o} + rt \tag{4.2}$$

Specific growth rate:

$$r(t) = \frac{d}{dt}\ln px_w(t) \tag{4.3}$$

$$r_{max} = \max_t\left(\frac{d}{dt}\ln px_w(t)\right) \tag{4.4}$$

Pixel-doubling time:

$$D = \frac{\ln(2)}{r_{max}} \tag{4.5}$$

### Method verification

The chosen method involves observing and evaluating the growth of the grains in three dimensions in a two-dimensional way. Therefore, a method evaluation was carried out to confirm the method's basic suitability. All calculations and statistical analyses were carried out at a significance level of 0.05.

#### Static grain mass - white pixels correlation analysis

The aim was to assess whether the number of white pixels in two-dimensional (2D) images of individual kefir grains can be used to indirectly reflect their three-dimensional (3D) properties, e.g., the volume. As mass is easier to determine and is related to volume via density, mass was selected as the 3D reference value. The relationship between the area in the top view, represented by the number of white pixels, and the experimentally measured grain mass was investigated. For this purpose, single grains were selected based on the size range from small to extra-large. The grains were washed two times with sterile tap water, then blotted dry with a lint-free cloth to remove any excess water, before being weighed. Each grain was placed in an individual crystallisation dish, and an image was taken. This image was then converted into an 8-bit format, and the number of white pixels $px_w$, using a ROI that exactly encloses the grain, was determined as described above.

The data analysis was based on a geometric model that describes the grains as approximately spherical. As a further approximation, it is assumed that the density of the grains remains constant. The relationship between mass and observed area is derived using formulas for the volume of a sphere (radius cubed) and the area of the largest cross-section (radius squared). Using the general formula for calculating density, the following equation can be derived.

$$m = \rho * \frac{4}{3}\pi * \left(\sqrt{\frac{A}{\pi}}\right)^3 = \rho * \frac{4}{3}\pi * \frac{A^{\frac{3}{2}}}{\pi^{\frac{3}{2}}} = \rho * \frac{4}{3} * \frac{A^{\frac{3}{2}}}{\pi^{\frac{1}{2}}} \quad (5.1)$$

Assuming a spherical geometry and constant density, a non-linear relationship between the visible area, represented by the number of white pixels, and the mass of a kefir grain is revealed.

$$m_{grain} \propto A_{pxw}^{\frac{3}{2}} \quad (5.2)$$

To test this relationship, a log-log regression using the natural logarithm $ln(x)$ was performed. The natural logarithms of the area, represented by the number of white pixels $ln(A_{pxw})$, and the corresponding measured mass of the individual kefir grains $ln(m_{grain})$, were calculated. A linear regression was performed with the number of white pixels as the independent variable and the grain mass as the dependent variable. This approach allows testing a power-law relationship of the form $ln(m) = a+k*ln(A)$, where $k$ corresponds to the slope of the regression line. Under the taken assumptions, an ideal slope of the log-log regression would be 1.5.

**Comparison of the dynamic 2D pixel number and the 3D mass increase**

In addition to comparing the instantaneous mass and number of white pixels generated by a single grain, the comparability of the increases in the measured parameters was also assessed. Previously, the grains were approximated as spherical. However, growth does not proceed uniformly in all spheres, and numerous irregularities can occur in the grain shape. To investigate this, a growth experiment was carried out as described above. The grains were washed twice with sterile tap water, then blotted dry with a sterile, lint-free cloth to remove any excess water before being weighed. Each grain was then placed in a separate crystallisation dish and observed for 72 h. After the last image was taken, the grains were taken out, blotted dry with a lint-free cloth, and weighed again.

For the data analysis, the number of white pixels in each grain was first determined in the initial image and in the final image after 72 h. To achieve this, a specific ROI was used for each image and grain to precisely enclose each grain, enabling comparison of the white pixels of the grains exclusively with the mass. The percentage increases in mass and white pixels, and the arithmetic means were then calculated from the respective start and end values. Before further analysis, the individual values were checked for outliers using the Dixon test. To finally compare the two methods, a paired-sample t-test was performed, as the measurements from both methods were taken from the same samples.

### Supplemental results

#### Exopolysaccharide production of kefir isolates on MRS agar supplemented with sucrose

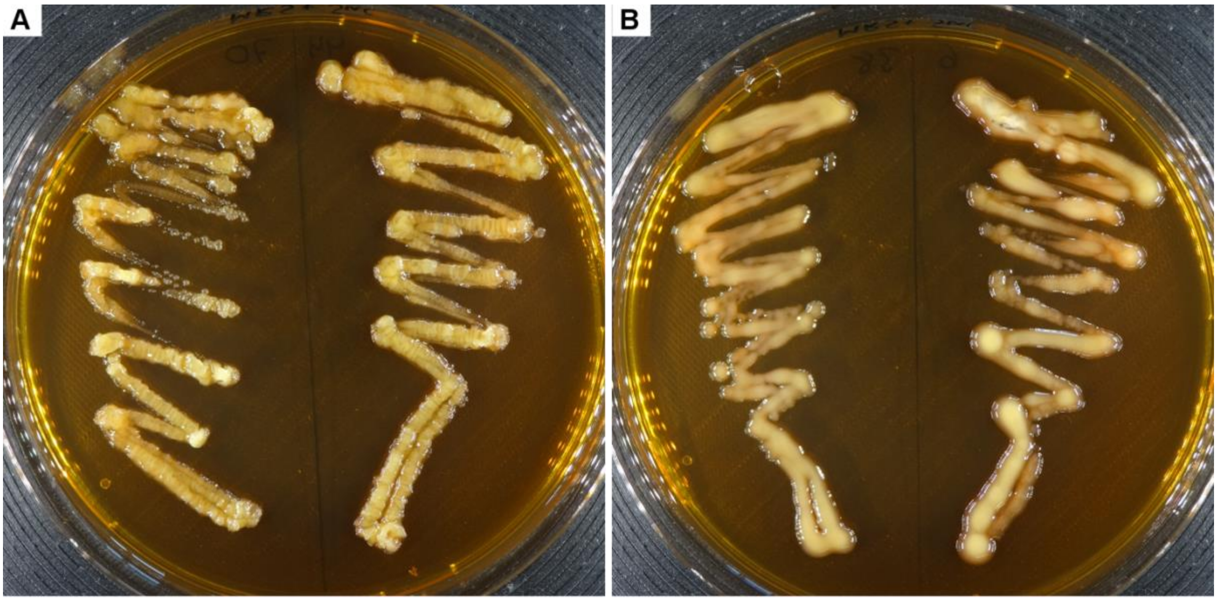


Figure S3. Exopolysaccharide production of kefir isolates on De Man, Rogosa, and Sharpe (MRS) agar supplemented with 5% (wt/vol) sucrose. (**A**) *Liquorilactobacillus satsumensis*; left: isolate TMWK070, right: isolate TMWK044. (**B**) *Liquorilactobacillus nagelii*; left: isolate TMWK038, right: isolate TMWK006. Agar plates were documented using a P.CAM360 with default light settings.

### Water kefir similarity network

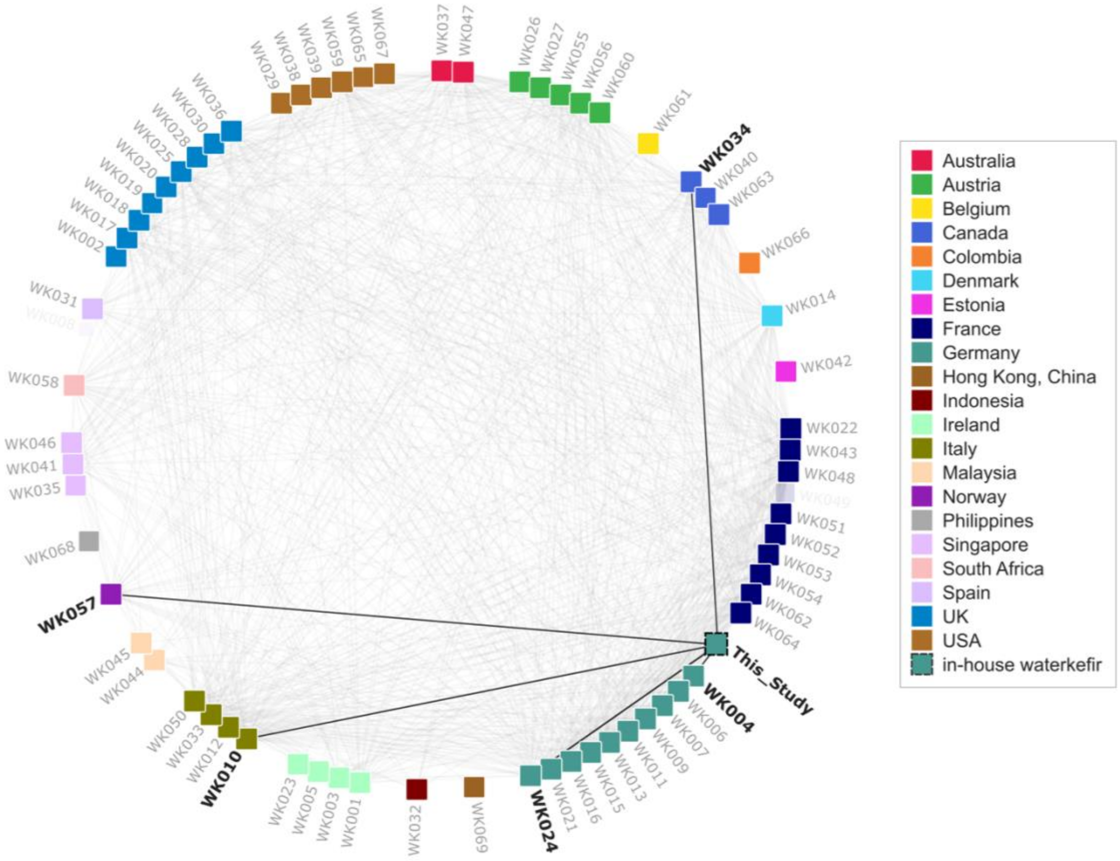


Figure S4. Sample–sample similarity network of water kefir communities based on species-level Jaccard similarity. Nodes represent individual water kefir samples, including the culture analysed in this study and the 69 reference samples reported by [32]. Edges connect samples with a conservative Jaccard similarity of at least 0.25, calculated from species-level presence–absence profiles. The five edges connecting the culture analysed in this study to its most similar reference samples are highlighted by increased line width, while edges between reference samples are shown with lower visual emphasis. Four of these five samples were also among the five nearest reference samples under the relaxed Jaccard comparison, in which ambiguous assignments were evaluated based on overlap with their possible species identities; WK024 remained the single most similar sample under both metrics. The conservative Jaccard similarities for the five closest reference samples highlighted in this figure ranged from 0.25 for WK004 to 0.33 for WK024. The highest conservative Jaccard similarity, 0.76, was observed between WK015 and WK016.

### Supplemental data and plots on flow cytometry and amplicon sequencing of selected subcommunities

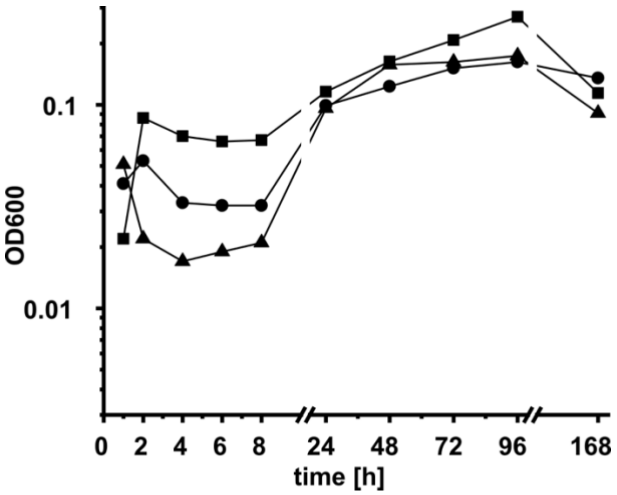


Figure S5. $OD_{600}$ measured in the fermentation experiment for subsequent cytometric analysis in batch replicate 1 (■), 2 (●), and 3 (▲).

Table S2. Fermentation process parameters pH, total soluble solids (TSS), and grain mass measured in the fermentation experiment for subsequent cytometric analysis. Data are presented as mean ± SD (*n* = 3).

| parameter | timepoint [h] | value |
|---|---|---|
| **pH** | 0 | 6.80 ± 0.0 |
| | 168 | 3.17 ± 0.02 |
| **TSS [°Brix]** | 0 | 7.8 ± 0.0 |
| | 168 | 1.6 ± 1.9 |
| **grain mass [g]** | 0 | 50.0 ± 0.0 |
| | 168 | 95.4 ± 6.4 |

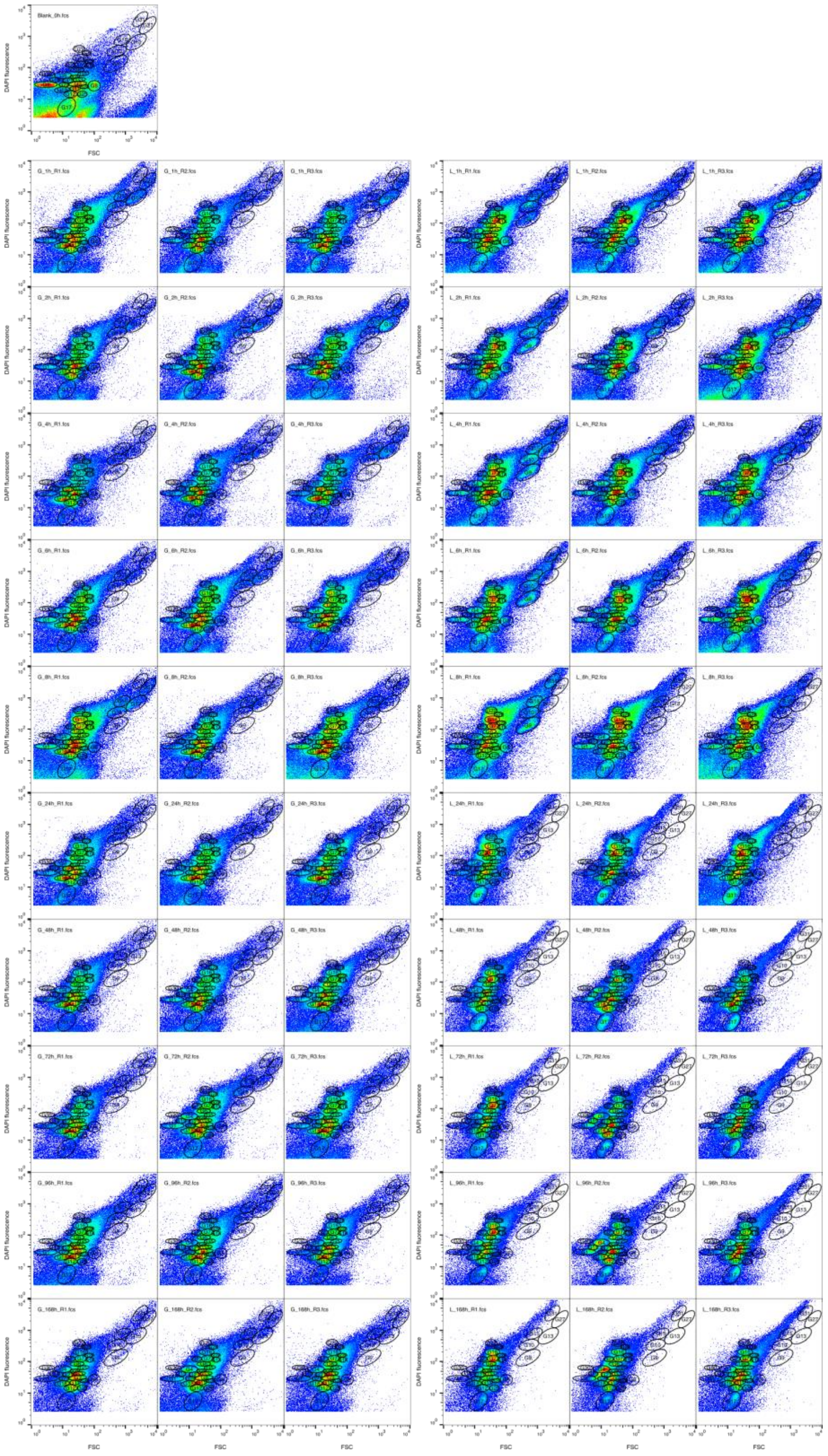

Figure S6. Overview of 2D flow cytometric plots of DAPI fluorescence intensity versus forward scatter (FSC) of grain- and liquid-associated microbial communities during water kefir fermentation. The blank consisted of the fermentation substrate containing sugar and dried fruit without water kefir grains. A uniform template comprising 31 numbered gates was applied to all samples and is shown in Figure 3. Sample labels indicate the sample fraction (G, grain; L, liquid), sampling time in hours after the start of fermentation, and replicate number (R1–R3).

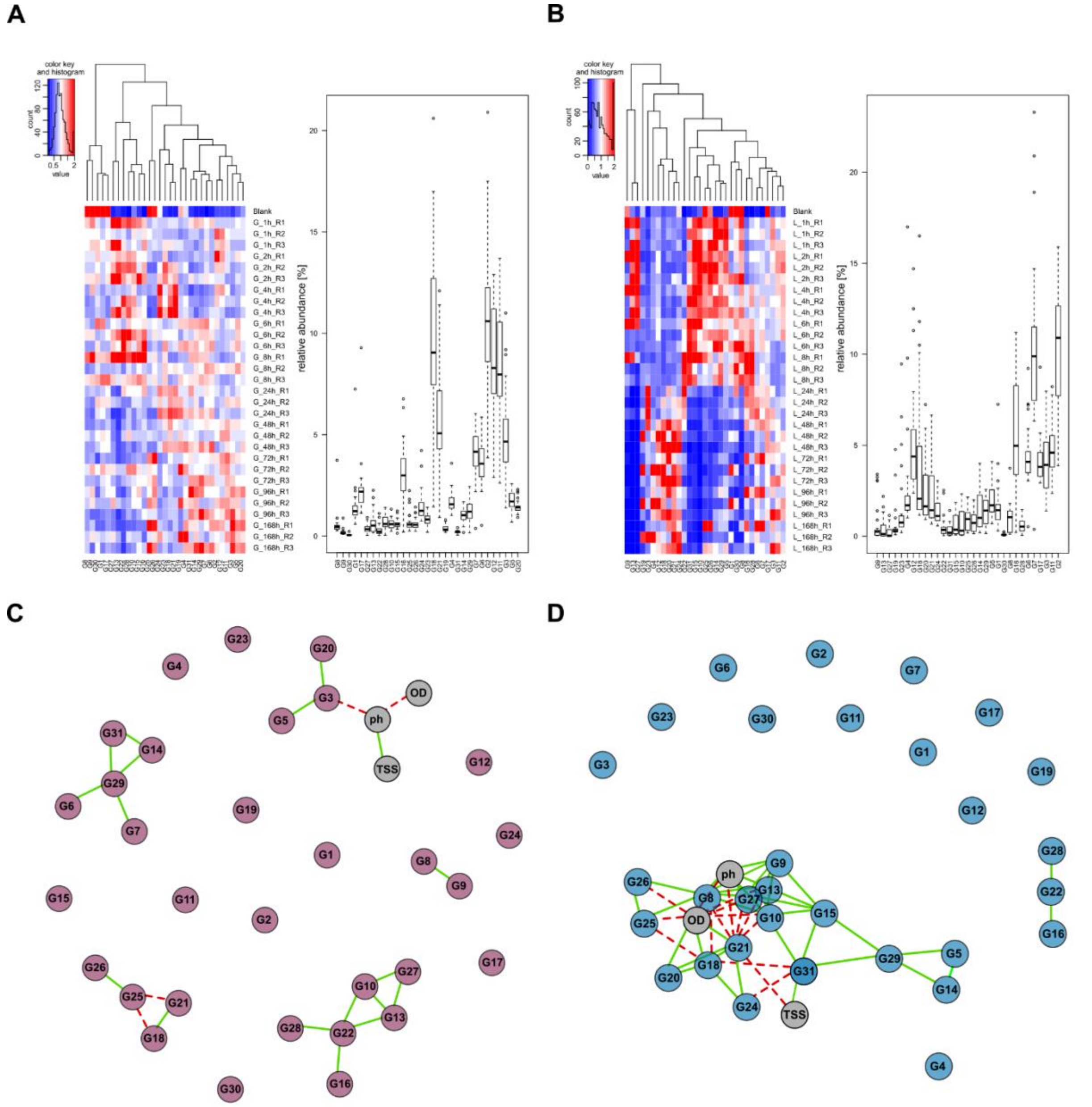


Figure S7. Cytometric gate dynamics and correlation analyses for grain and liquid fractions. (**A–B**) Heat maps displaying the variations in relative gate abundances over time and across three biological replicates for grains (**A**) and liquid (**B**). Gates are ordered by similarity. Corresponding box plots indicate overall gate abundances across all samples; outliers are marked by circles. (**C–D**) Spearman's rank order correlation maps ($r \geq 0.75$, $p \leq 0.05$) for grains (**C**) and liquid (**D**), showing significant associations between gates and the selected process parameters $OD_{600}$, pH, and TSS. Positive correlations are indicated by solid green lines, negative correlations by dashed red lines.

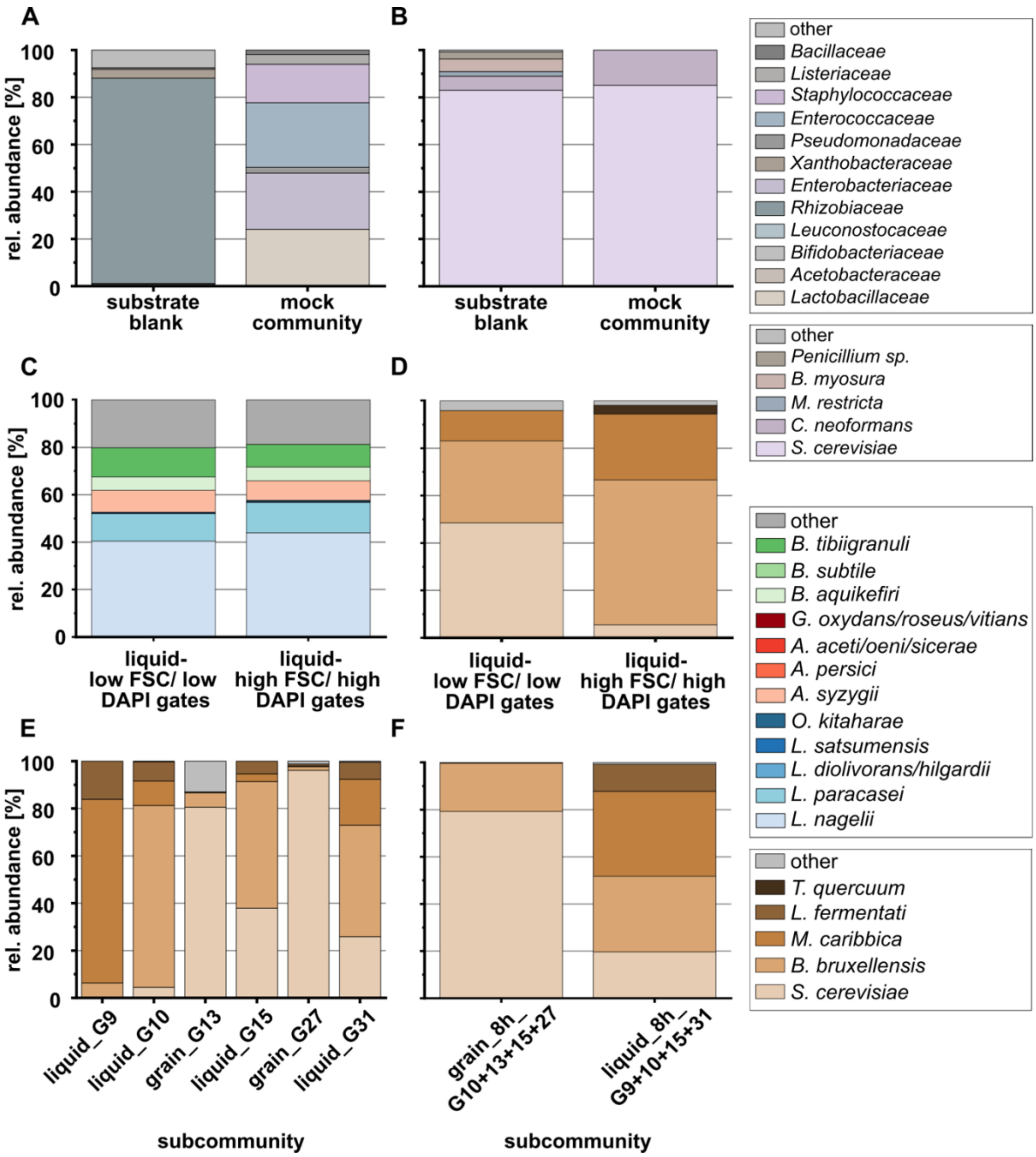


Figure S8. Taxonomic composition of selected FACS-sorted subcommunities based on ASV-resolved 16S and ITS amplicon sequencing. Panel A is shown at the family level for clarity; all other panels depict species-level taxonomic resolution. Combined species names indicate that the respective sequence could not be reliably resolved to a single species based on the amplified marker region. (**A**) 16S rRNA gene-based community profiles of the substrate blank and the ZymoBIOMICS mock community and (**B**) ITS-based profiles of the same samples. (**C–D**) Taxonomic composition of combined gates from the liquid fraction, representing low FSC/low DAPI fluorescence and high FSC/high DAPI fluorescence regions, based on (**C**) 16S and (**D**) ITS amplicon sequencing, respectively. Samples were taken from batch replicate 1 from the liquid after 4 h of fermentation. (**E**) ITS-based community profiles of individual high FSC/high DAPI gates, sorted as putative yeast-enriched subcommunities from grain or liquid samples. (**F**) ITS-based profiles of pooled high-abundance high FSC/high DAPI fluorescence gates sorted after 8 h of fermentation in grains and liquid (four gates each).

### Two-dimensional image analysis reflects three-dimensional grain size and growth

To evaluate the validity of using 2D image data as a proxy for 3D grain properties, the relationship between image-derived pixel counts and the mass of individual kefir grains was examined, with mass serving as a convenient proxy for volume. In the first experiment, kefir grains ranging from 2 mm (small) to 10 mm (extra-large) were analysed by recording their mass and white pixel counts. Log-log regression, based on a geometric model of spherical shape and constant density, yielded a slope of 1.35 ± 0.11 with $R^2 = 0.98$ (Figure S9), slightly lower than the theoretical value of 1.5. This indicates that mass increases less than proportionately with pixel counts, particularly in large grains, due to more frequent structural irregularities such as cavities or indentations.

To extend the analysis beyond static measurements, a second experiment monitored the growth of individual grains over 72 h. The relative mass increase (Table S3) was compared with the corresponding increase in white pixels (Table S4) to evaluate the consistency of the imaging approach under dynamic conditions. One sample deviated more strongly because the grain broke into two parts, resulting in a disproportionate increase in pixel counts relative to mass. Despite this, no outliers were detected, and no significant differences were found between the mean relative increases in mass and pixels ($p \leq 0.05$).

Together, both experiments confirmed a consistent geometric relationship between pixel counts and grain mass, validating the method for dynamic growth. White pixel counts thus provide a quantitative metric for growth, capturing relative differences and trends across grains from 2–10 mm in diameter, but not absolute volumetric values.

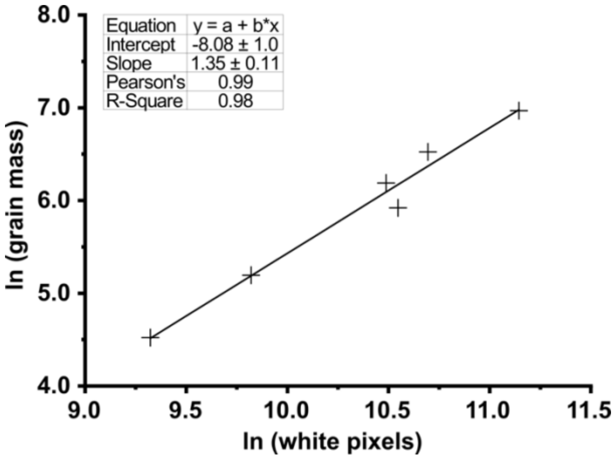


Figure S9. Regression curve and parameters for log-log regression of white pixels and single grain mass.

Table S3. Kefir grain mass of single grains at the beginning of the experiment and after 72 h, the calculated difference, and the relative increase for pairwise comparison of 2D- and 3D-properties.

| grain mass [g] 0 h | grain mass [g] 72 h | ∆ grain mass [g] | rel. increase [%] |
|---|---|---|---|
| **231.2** | 463.9 | 232.7 | 100.6 |
| **271.6** | 548.5 | 276.9 | 101.9 |
| **269.5** | 518.6 | 249.1 | 92.4 |

Table S4. Absolute numbers of white pixels from single kefir grains at the beginning of the experiment and after 72 h, the calculated difference, and the relative increase for pairwise comparison of 2D- and 3D-properties.

| white pixels [px] 0 h | white pixels [px] 72 h | ∆ white pixels [px] | rel. increase [%] |
|---|---|---|---|
| **20,426** | 41,316 | 20,890 | 102.3 |
| **20,924** | 43,617 | 22,693 | 108.5 |
| **22,134** | 47,898 | 25,764 | 116.4 |

### Supplemental kefir growth images, growth curves, and parameters for growth quantification

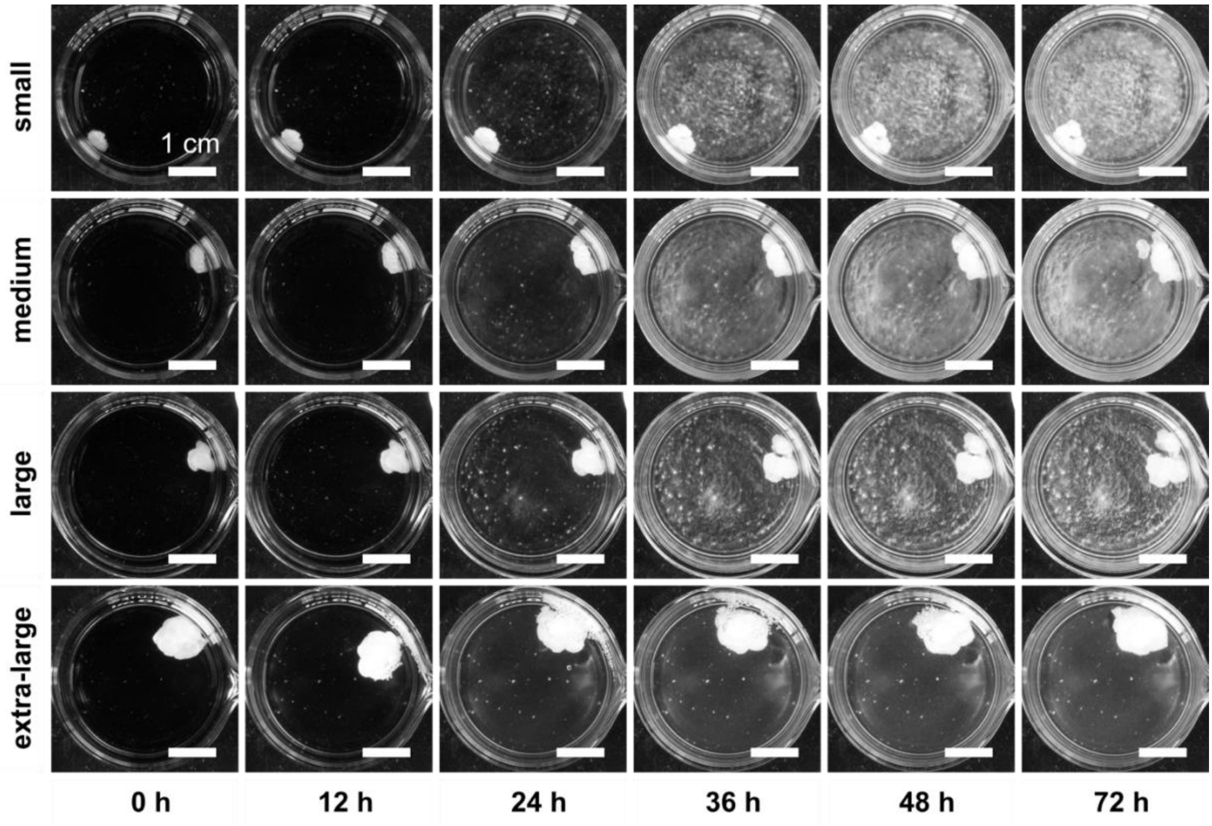


Figure S10. Representative 8-bit images illustrating the macroscopic growth behaviour of single water kefir grains with different initial sizes. Small, medium-sized, large, and extra-large starter grains were inoculated individually in water kefir medium and monitored over a 72-h observation period. The images correspond to the growth curves shown in Figure S13. Scale bars represent 1 cm.

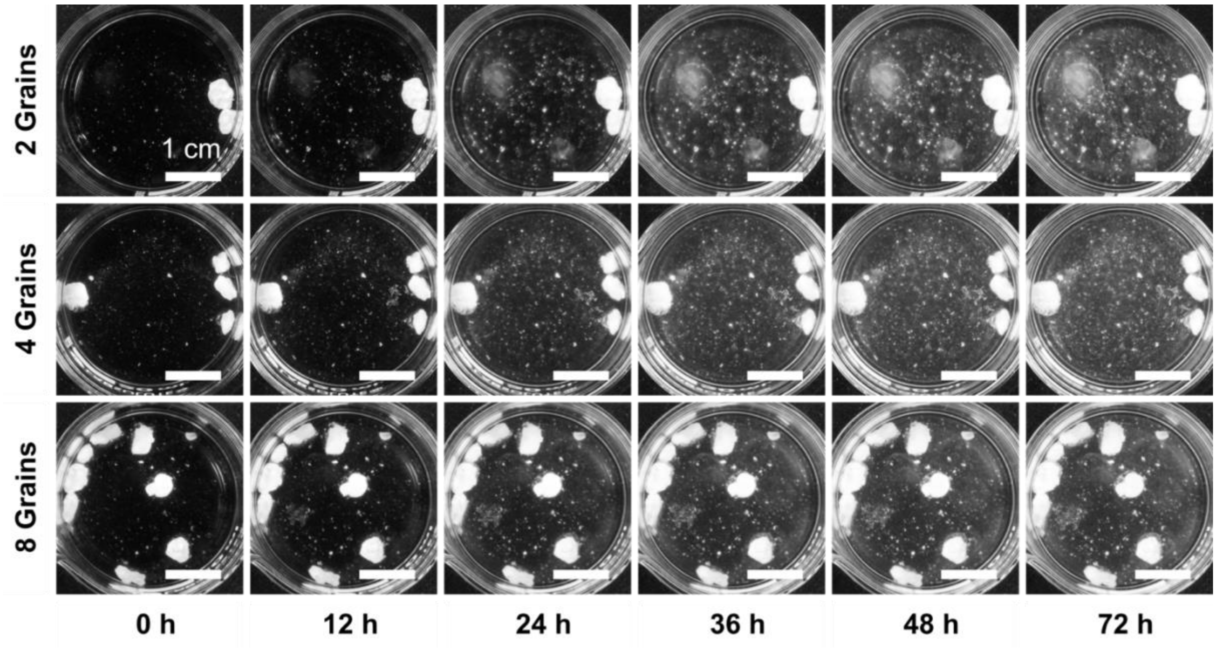


Figure S11. Representative 8-bit images illustrating the macroscopic growth behaviour of water kefir grains at different initial grain numbers. Experiments were conducted with 2, 4, and 8 medium-sized starter grains per vessel in water kefir medium, and the cultures were monitored over a 72-h observation period. Scale bars represent 1 cm.

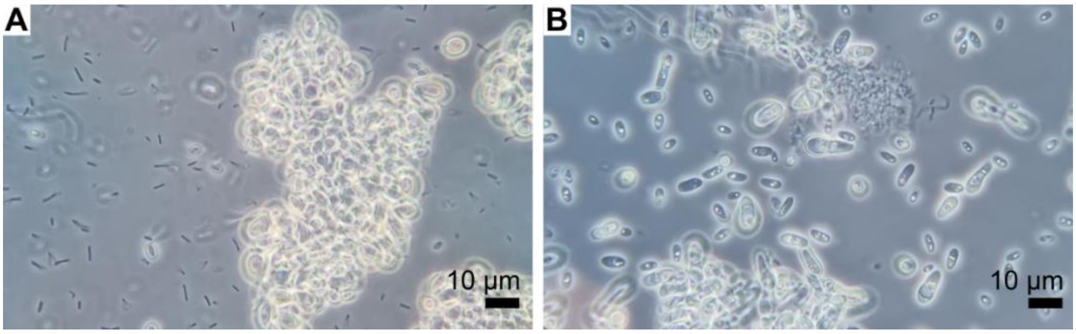


Figure S12. Microscopic images of (**A**) the water kefir liquid after 24 h of fermentation and (**B**) the biofilm at the bottom of the sample vessels from camera-based growth experiments. Microscopic images were acquired using phase-contrast microscopy at 1000x magnification.

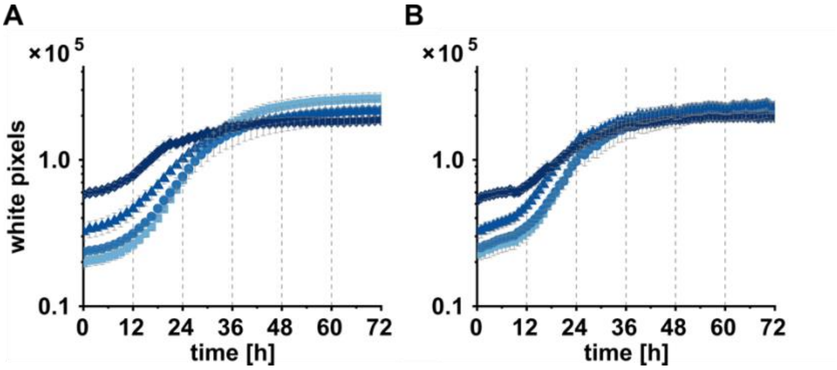


Figure S13. Growth curves of individual kefir grains of different initial sizes. Growth dynamics based on total ROIs for samples containing single kefir grains of different initial sizes: small (■), medium (●), large (▲), and extra-large (◆). Darker colours indicate larger starter grain size. (**A**) Experiment 1; (**B**) Experiment 2.

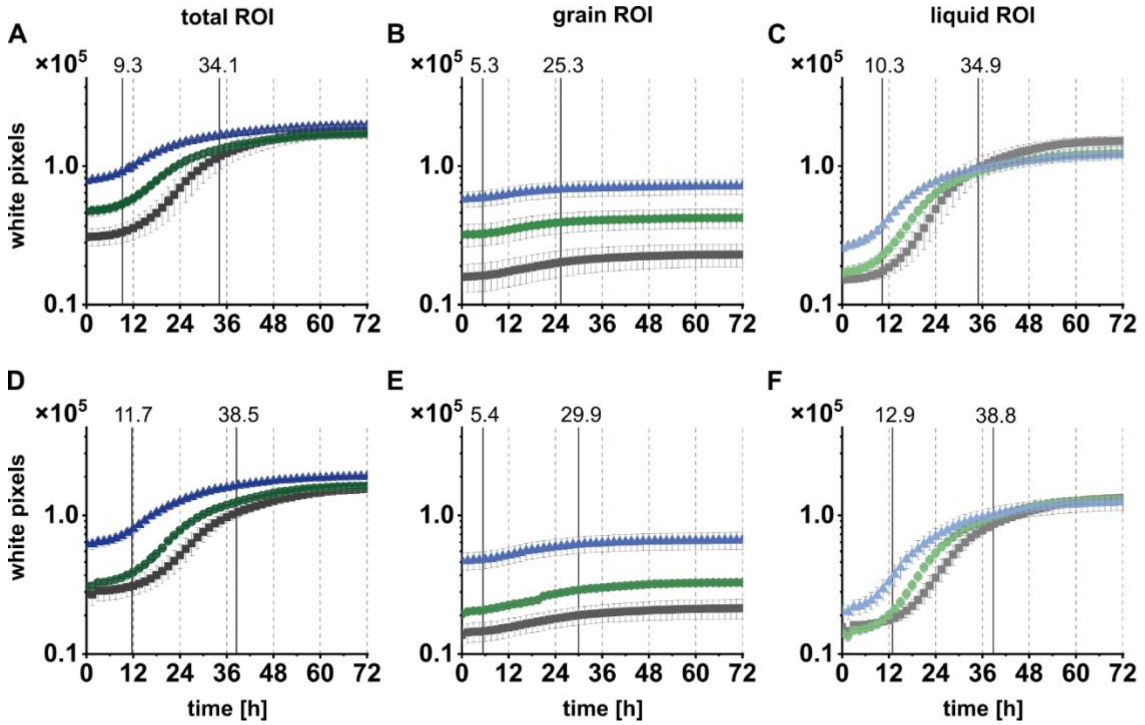


Figure S14. Growth curves of medium-sized kefir grains at different initial numbers for two experimental replication runs. (**A–C**) Experiment 2; (**D–F**) Experiment 3. Growth dynamics are shown for samples inoculated with 2 (■), 4 (●), and 8 (▲) medium-sized grains. Curves represent the mean values of three biological replicates (*n* = 3) analysed using different regions of interest: (**A, D**) total ROI, (**B, E)** grain ROI, and (**C, F**) liquid ROI. Vertical solid lines indicate the median time points at which 5% and 70% of the absolute increase in white pixels was reached across all nine samples in one experiment.

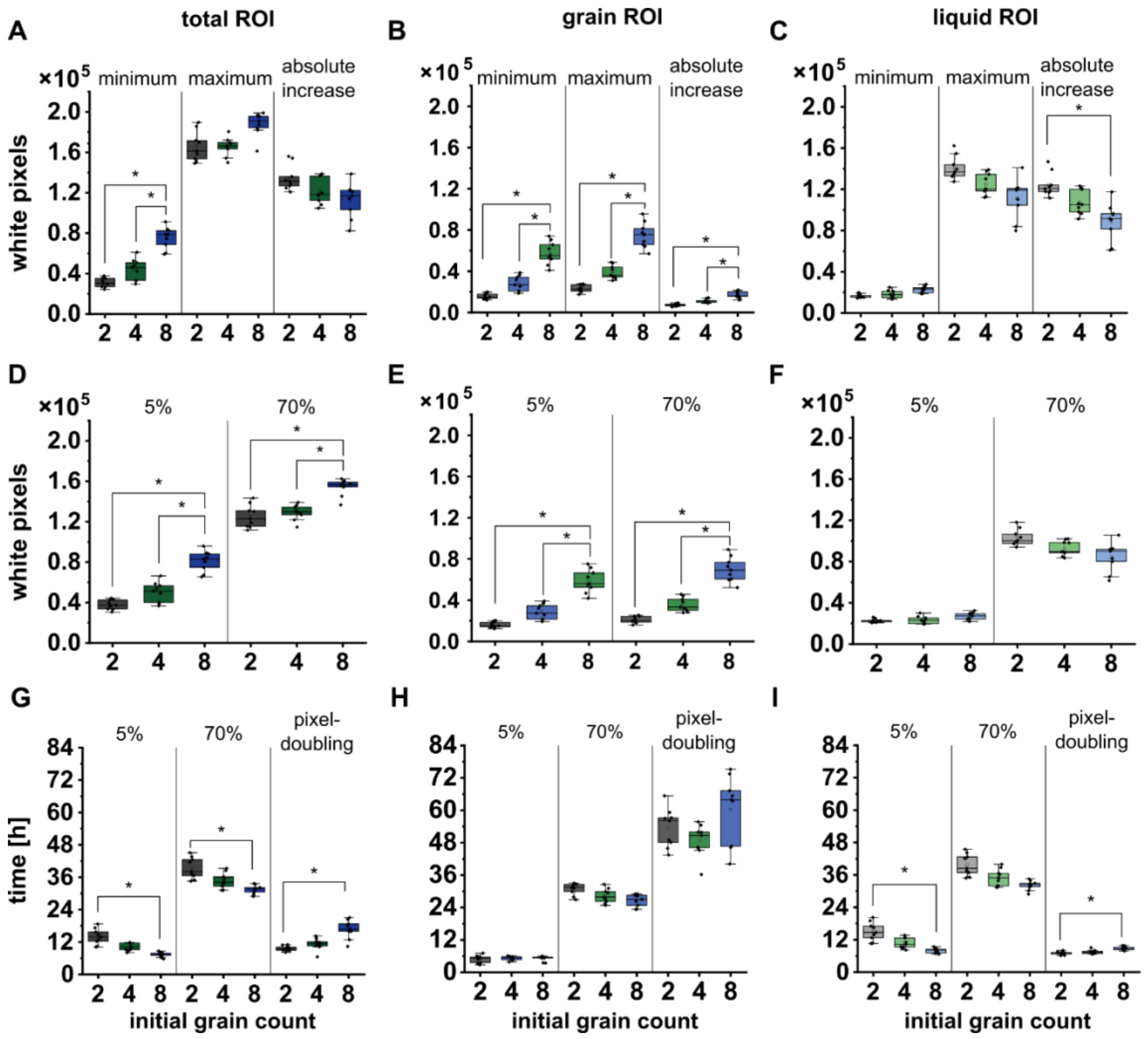


Figure S15. Box plots of growth parameters derived from different regions of interest for samples containing 2, 4, and 8 medium-sized starter grains. Panels show values derived from the total ROI (**A, D, G**), grain ROI (**B, E, H**), and liquid ROI (**C, F, I**). (**A–C**) Minimum, maximum, and absolute increase in white pixel counts. (**D–F**) White pixel counts at which 5% and 70% of the absolute increase were reached. (**G–I**) Time required to reach 5% and 70% of the absolute increase, and the calculated pixel-doubling time. Box plots summarise all measurements from three independent 72-h experiments, while statistical analyses were performed on the means of three biological replicates per group (one-way ANOVA with Tukey's post hoc test, $n = 3$). Asterisks indicate significant differences ($p \leq 0.05$).

Across the total ROIs, which captured all ongoing growth processes, the minimum pixel values increased with grain number, while maximum values remained within a comparable range (Figure S15A). Consequently, the absolute increase tended to be lower at higher inoculation volumes. For both thresholds, the pixel values increased with the initial grain number (Figure S15D), while the timepoints at which the thresholds were reached showed an inverse trend (Figure S15G). As the inoculum size increased, both thresholds were reached earlier, as a smaller difference in white pixels had to be overcome between the lower and upper thresholds. The lower threshold was reached after 9–12 h. This was followed by the exponential growth phase, with the curves reaching the upper threshold after 32–39 h and subsequently entering a stationary phase without decline. The estimated pixel-doubling times increased with the initial grain number, indicating slower relative growth at higher biomass inputs (Figure S15G).

The same approach was used for the grain sub-ROIs to further distinguish the contributions of the individual phases to the overall growth behaviour. The minimum, maximum, and total pixel values increased with the number of grains (Figure S15B), and the same pattern was observed for pixel values at the lower and upper thresholds (Figure S15E). A shorter lag phase and earlier saturation of the grain-associated area relative to the overall growth were indicated by the lower threshold being reached after 5–6 h and the upper threshold after 25–30 h (Figure S15H). However, there were no significant differences in timing parameters or pixel-doubling times between sample groups. The white-pixel data for the grains generally showed greater variability, as slight grain movements influenced measurements more strongly in this grain sub-ROI than in the total ROI.

The growth curves derived from the liquid sub-ROIs showed lower absolute pixel values and a more pronounced sigmoidal shape than those derived from the total ROI. In general, the timing of growth in the liquid phase was comparable to total growth but delayed compared to grain growth. As the grain number per vessel increased, the available liquid ROI area and thus the number of possible pixel values decreased. Parameter analysis confirmed that liquid ROI data were significantly affected by spatial constraints and could not be interpreted independently of one another. Minimum pixel values were nearly identical across conditions due to the initially uncolonized liquid, while maximum values and the absolute increase decreased with higher grain numbers (Figure S15C). Although the pixel values at growth thresholds were similar between conditions (Figure S15F), the time points varied. Both thresholds were reached earlier at higher grain numbers, suggesting faster initial and general activity (Figure S15I). The lower threshold was reached after 9–13 h, followed by an exponential increase in pixel intensity due to turbidity and biofilm formation. The upper threshold was reached between 32 and 39 h. However, pixel-doubling time increased with inoculum size, reflecting spatial and nutrient limitations in samples with large initial biomass (Figure S15I).

### Supplemental images on kefir grain morphology

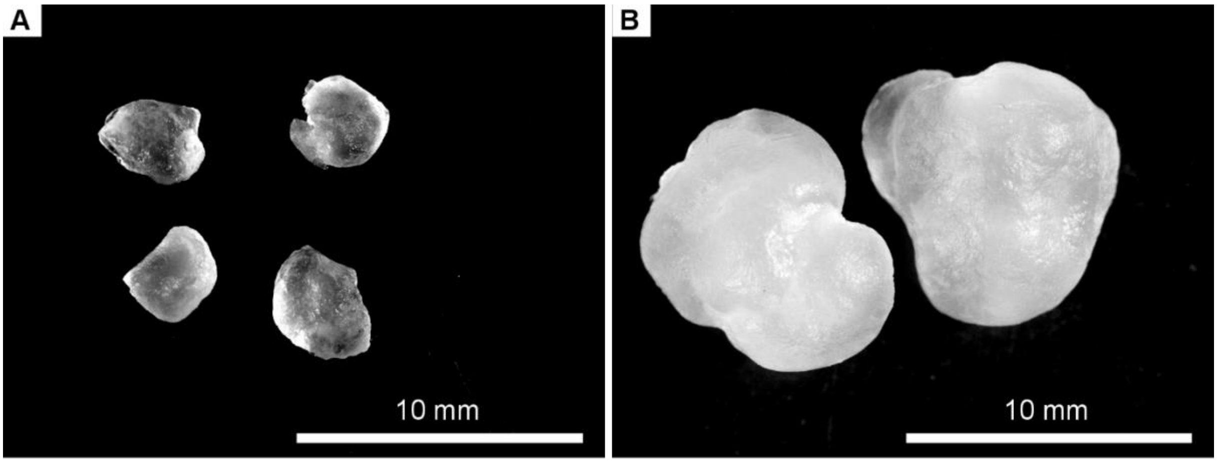


Figure S16. Macroscopic morphology of water kefir grains of different sizes: (**A**) small, (**B**) large to extra-large. Stereomicroscopic images are shown in grayscale and were acquired at 10x magnification.
